# Metabolism regulates muscle stem cell self-renewal by connecting the microenvironment and histone acetylation

**DOI:** 10.1101/2023.07.04.547746

**Authors:** C. Hai Ly, Jin D. Chung, John H.V. Nguyen, Luyi Tian, Jan Schroeder, Anja S. Knaupp, Shian Su, Jennifer Trieu, Talhah M. Salmi, Daniela Zalcenstein, Jafar S. Jabbari, Berin A. Boughton, Andrew G. Cox, Shalin H. Naik, Jose M. Polo, Matthew E. Ritchie, Gordon S. Lynch, James G. Ryall

**Affiliations:** Centre for Muscle Research, Department of Anatomy and Physiology, The University of Melbourne, Victoria, 3010, Australia; The Walter and Eliza Hall Institute of Medical Research and Department of Medical Biology, The University of Melbourne, Victoria, 3010, Australia; Australian Regenerative Medicine Institute, Monash University, Wellington Road, Clayton, Victoria, 3800, Australia; Cancer Metabolism Program, Organogenesis and Cancer Program, Peter MacCallum Cancer Centre, Melbourne, Victoria, Australia; Australian Genome Research Facility, Melbourne, Victoria, Australia; Metabolomics Australia, The University of Melbourne, Melbourne, Victoria 3010, Australia; Australian National Phenome Center, Murdoch University, Murdoch, Western Australia 6150, Australia

**Keywords:** citrate, glycolysis, myogenesis, satellite cell, asymmetric division, scRNAseq

## Abstract

Skeletal muscle contains a resident population of somatic stem cells capable of both self-renewal and differentiation. The signals that regulate this important decision have yet to be fully elucidated. Here we use metabolomics and mass spectrometry imaging (MSI) to identity a state of localized hyperglycaemia following skeletal muscle injury. We show that committed muscle progenitor cells exhibit an enrichment of glycolytic and TCA cycle genes and that extracellular monosaccharide availability regulates intracellular citrate levels and global histone acetylation. Muscle stem cells exposed to a reduced (or altered) monosaccharide environment demonstrate reduced global histone acetylation and transcription of myogenic determination factors (including *myod1*). Importantly, reduced monosaccharide availability was linked directly to increased rates of asymmetric division and muscle stem cell self-renewal in regenerating skeletal muscle. Our results reveal an important role for the extracellular metabolic environment in the decision to undergo self-renewal or myogenic commitment during skeletal muscle regeneration.

## INTRODUCTION

Skeletal muscle contains a resident population of somatic stem cells (muscle stem cells or MuSCs) that confer a high regenerative capacity on this tissue. MuSCs exist in a quiescent state outside of the cell cycle and are marked by the paired box transcription factor Pax7. In response to an activating signal, MuSCs rapidly exit quiescence, enter the cell cycle and undergo several rounds of proliferation, a period that coincides with the appearance of the bHLH transcription factors, Myf5 and MyoD, and commitment to the myogenic lineage (Hernandez-Hernandez et al., 2017; Zammit, 2017). Importantly a subpopulation of MuSCs do not undergo commitment and instead return to quiescence. This self-renewing property of MuSCs prevents depletion of the stem cell pool and maintains the regenerative capacity of skeletal muscle. Any change to this balance of self-renewal versus commitment can have devastating consequences on regenerative capacity (Chang et al., 2018; Dumont et al., 2015).

While extensive work over the last 30 years has helped define (and refine) the transcriptional network that regulates MuSC activation, specification, and differentiation (Fukada et al., 2007; Machado et al., 2017; Pallafacchina et al., 2010), we are only now beginning to understand the cellular signals that regulate these complex transcriptional pathways. Cellular signals can include biophysical cues, autocrine and paracrine signaling, and environmental stimuli such as the pH and pO_2_ of the local tissue environment. More recently, innate cell metabolism has been identified as a key regulator of MuSC biology, with quiescent MuSCs exhibiting an enrichment of fatty acid oxidation genes and activated/proliferating MuSCs switching to a highly glycolytic phenotype (Machado et al., 2017; Ryall et al., 2015b). Proliferating cells are typically reliant on a steady supply of the monosaccharide glucose for both ATP and new biomass, in the form of nucleotides, amino acids, and phospholipids (Hosios et al., 2016; Lunt and Vander Heiden, 2011). However, in addition to meeting the bioenergetic demands of dividing cells, the extracellular environment plays an important role in the regulation of both lineage progression and the epigenome (Ly et al., 2020; Ryall et al., 2015a). For example, reduced fatty-acid oxidation impairs the activation of MuSCs while driving precocious differentiation (Gatta et al., 2017; Pala et al., 2018). In addition, the NAD-dependent SIRT1 maintains histone H4K16 in a deacetylated state in quiescent MuSCs (when [NAD] is highest) leading to transcriptional repression (Ryall et al., 2015b). However, depletion of NAD and the subsequent decline in SIRT1 mediated deacetylation alone is not sufficient to explain the histone acetylation observed during MuSC activation.

The tricarboxylic acid (TCA) cycle is central for the production of several key metabolites that modulate the epigenome, including citrate. Citrate can be actively transported out of the mitochondria and converted to acetyl-CoA, which can then diffuse across the nuclear membrane and interact with histone acetyltransferases (Cai et al., 2011; Wellen et al., 2009). Therefore, in addition to regulating [NAD], increased glycolytic flux feeding the TCA cycle may directly promote histone acetylation and provide a temporal link between the extracellular environment and the regulation of transcription.

Here we show that perturbations in skeletal muscle architecture are accompanied by localized hyperglycemia leading to remodeling of the local tissue metabolic microenvironment. We demonstrate that in proliferating MuSCs, extracellular monosaccharide availability regulates cell fate. Our results suggest that in conditions of reduced substrate availability, activated MuSCs prioritize the production of biomass and cell division over the supply of pyruvate to the TCA cycle. To meet ongoing energetic demands, MuSCs divert citrate/acetyl-CoA from histone acetylation to maintain TCA cycle anaplerosis. Interestingly, under these conditions of reduced histone acetylation, MuSCs preferentially undergo asymmetric division, leading to increased Pax7^+^ and MyoD^-^ MuSCs at the expense of Pax7^-^ and MyoD^+^ muscle progenitors. Our results directly implicate the local metabolic milieu in the temporal regulation of transcription and the process of myogenic lineage progression.

## RESULTS

### Metabolomic analysis of regenerating skeletal muscle reveals local hyperglycemia at the site of injury

We sought to generate a metabolomic signature of skeletal muscle repair during a period of rapid cell proliferation and one marked by differentiation and maturation (Figure 1A). To study muscle regeneration, we injured the right tibialis anterior (TA) hindlimb muscles of C57BL/6 mice with the myotoxic agent BaCl_2_, and then rapidly excised the muscles at 3- and 7-days post injury (dpi). Polar metabolomic analysis with targeted liquid chromatography-tandem mass spectrometry (LC/MS-MS) was then performed on control uninjured muscles, at 3 dpi and 7 dpi, as described previously (Cox et al., 2018). A 2D principal component analysis (explaining over 70% of variance) confirmed three distinct clusters for control, 3 dpi and 7 dpi samples (Figure 1B), while hierarchical clustering of metabolite abundance revealed distinct metabolite signatures for 3 dpi and 7 dpi compared to uninjured skeletal muscles (Figure 1C and Table S1).

**Figure 1.**
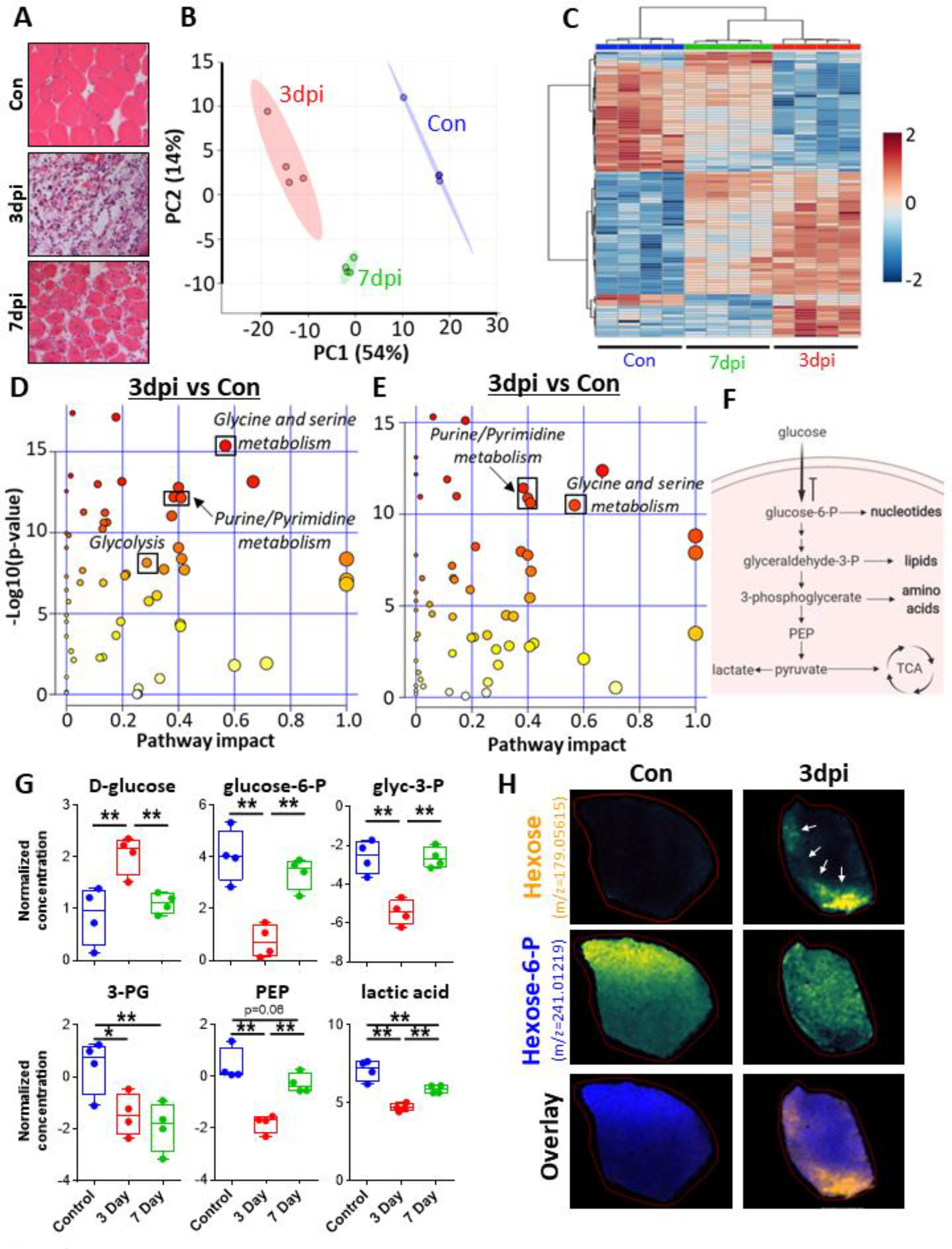
Early Skeletal Muscle Regeneration is Linked to Local Hyperglycemia. **(A)** Hematoxylin and eosin staining of TA muscles from control uninjured mice, and 3- or 7- days post-BaCl_2_-induced injury. (B) Principal component analysis of metabolomics performed on TA muscles from control uninjured mice or 3- or 7-days post-BaCl_2_-induced injury (n=4/timepoint). (C) Clustered heat-map of differential metabolite expression over the 7-day regeneration timeline (n=4/group). (D-E) Pathway enrichment analyses for differentially expressed metabolites at day 3 (D), or day 7 (E) of regeneration compared to uninjured control muscles. (F) Schematic outlining key glycolytic metabolites and their contributions to the production of new biomass. (G) Relative intramuscular concentration of six key glycolytic metabolites following BaCl_2_- induced injury (n=4/timepoint). (H) Matrix-assisted laser desorption/ionization–mass spectrometry imaging (MALDI-MSI) of hexose and hexose-6-P in TA muscles from control uninjured mice, or 3-days post submaximal BaCl_2_-induced injury. Arrows indicate regions of damaged tissue. Images were RMS normalized and viridis colored with signal intensity scaled linearly from 0-100%.

Skeletal muscle injury was linked with differential concentrations of metabolites involved in the synthesis of new biomass, including *glycine and serine metabolism*, *purine metabolism*, and *pyrimidine metabolism* (Figure 1D,E). Interestingly, the steady-state levels of metabolites involved in the central glycolytic pathway (Figure 1F) were differentially regulated at 3 dpi, but not at 7 dpi (Figure 1D,E). Specifically, D-glucose was elevated at 3 dpi compared to control muscles (Figure 1G). In contrast, the glycolytic intermediates, glucose-6-phosphate (G6P), glyceraldehyde-3-phosphate (G3P), 3-phosphoglycerate (3PG), phosphoenolpyruvate (PEP) and lactic acid, were all reduced at 3 dpi. By 7 dpi, the glycolytic intermediates upstream of 3PG had all returned to control levels (Figure 1G).

To better understand the observed local hyperglycemia at the site of muscle injury we switched to matrix-assisted laser desorption/ionization (MALDI) mass spectrometry imaging (MSI). Using a submaximal BaCl_2_ injury model, we imaged control uninjured TA muscles and TA muscles at 3 dpi. Similar to our metabolomics results, discriminative MSI analysis using Receiver Operating Characteristic (ROC) Area Under the Curve (AUC) of the overall MSI images revealed that hexose sugar was strongly discriminant for the injured TA muscle sample (AUC 0.138) and was highly localized to the injured TA tissues, while hexose-6-P was enriched in healthy uninjured tissue (Figure 1H), (AUC 0.869). This observed local hyperglycemia at 3 dpi coincides with a period of rapid cell proliferation prior to the formation of centrally nucleated myofibers (Figure 1A)(Hardy et al., 2016), and is likely important for the production of new biomass critical for dividing cells (Hosios et al., 2015; Nguyen et al., 2019).

### Single cell RNA sequencing of MuSCs isolated from 3 dpi regenerating skeletal muscles reveals distinct sub-populations of cells that exhibit divergent metabolic signatures

Several studies have characterized the metabolic signature of quiescent/freshly isolated and activated MuSCs using whole transcriptome sequencing (Machado et al., 2017; Ryall et al., 2015b). To determine if heterogeneity exists within either cell state, we performed single cell RNA sequencing (scRNAseq). To rapidly and efficiently isolate a large number of primary MuSCs we utilized Pax7^creERT2^R26-eYFP^fl/fl^ mice where tamoxifen administration leads to irreversible eYFP labelling of Pax7^+^ MuSCs (Figure S1A), which were then purified by fluorescence activated cell sorting (FACS, Figure S1B). To better understand the cell-specific signaling that may regulate the decision to either commit or self-renew following division, we performed scRNAseq on MuSCs freshly isolated from the uninjured TA muscles of male mice or the TA muscles of female mice at 3 dpi. All cells were then combined and barcoded libraries were prepared on a 10× chromium instrument before sequencing (Figure S1A). After normalization and quality control, we obtained 791 cells with a mean of >2.3 × 10^5^ reads/cell (mapping to over 16,00 total genes). Injured and uninjured cells were identified by the presence of sex-specific genes (*ddx3y* and *xist*, Figure S1C). Following cell identification, all sex-specific genes were removed and a two-dimensional t-distributed stochastic neighbor embedding (t-SNE) analysis (Figure 2A) with Seurat clustering was performed (Figure 2B).

**Figure 2.**
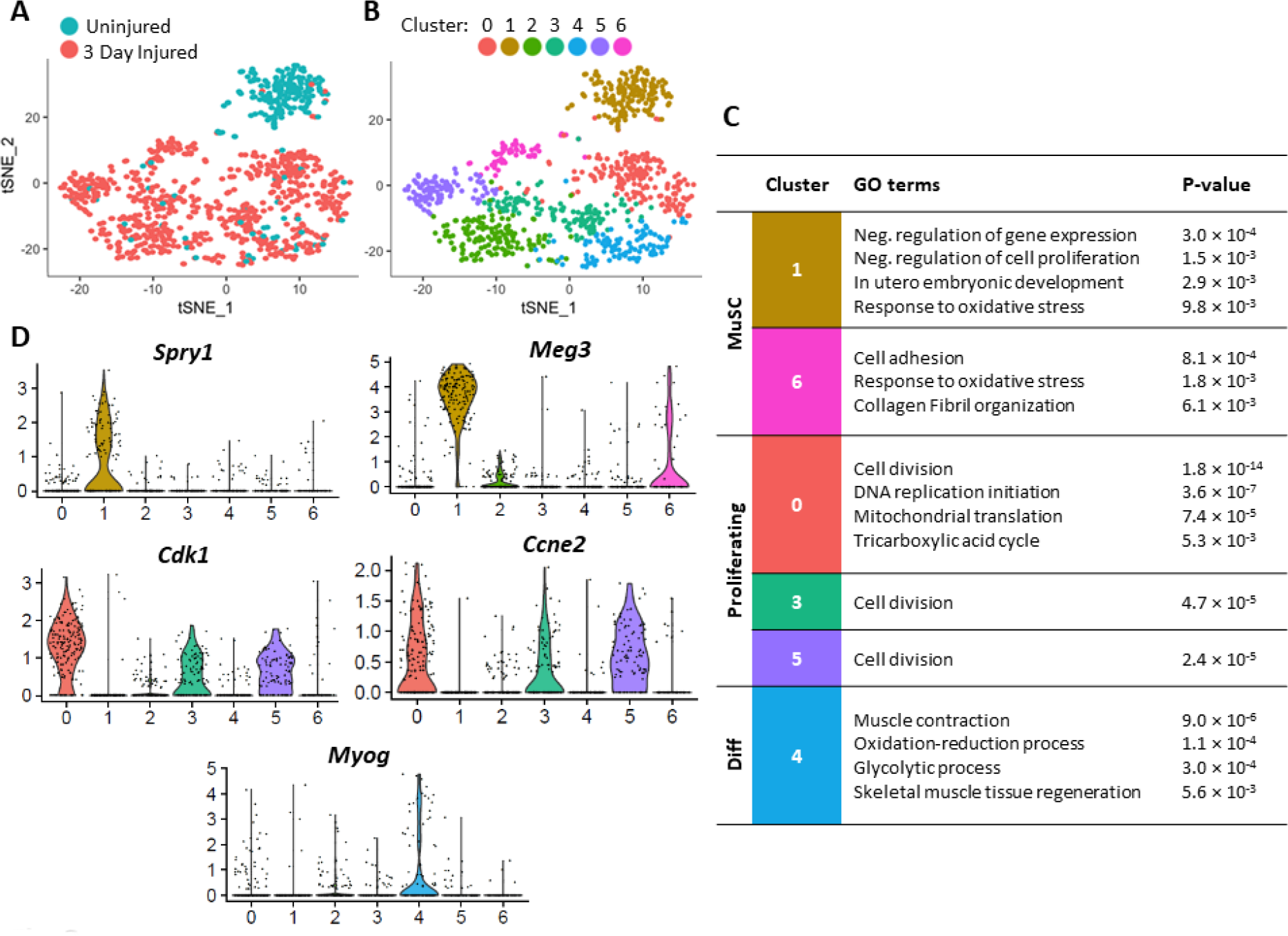
Single Cell RNAseq of MuSCs Freshly Isolated from Control and Injured Skeletal Muscle Reveals Distinct Innate Metabolic Profiles. (A) Single MuSCs freshly isolated from control and injured (3 dpi) skeletal muscles were mapped to two-dimensions using a tSNE approach (n=791 cells total) (B) Seven independent cell clusters were identified via Seurat. Cluster 1 was predominantly composed of MuSCs from uninjured muscles, while clusters 0 and 2-6 were composed primarily of MuSCs from injured muscles. (C) Gene ontology analyses of biological processes enriched in each cluster revealed distinct metabolic profiles. (D) Violin plots of gene expression for marker genes in clusters 1 and 6 (*Spry1 and Meg3*), clusters 0, 3 and 5 (*Cdk1 and Cccne2*) and cluster 4 (*Myog*).

Gene ontology (GO) analyses of biological processes enriched in clusters 1 and 6 revealed GO terms typically associated with quiescent MuSCs, including “*negative regulation of gene expression”, “negative regulation of cell proliferation”* and “*cell adhesion”* (Figure 2C). Similarly, an analysis of individual genes revealed an enrichment of several previously linked to a quiescent state, including *Spry1* and *Meg3* (Figure 2D and Table S2). While cluster 1 was made up of >95% of MuSCs isolated from uninjured muscles and therefore expected to be enriched for terms linked to quiescence, cells in cluster 6 primarily originated from 3 dpi muscles.

In contrast to clusters 1 and 6, clusters 0, 3 and 5 were all enriched for terms linked with an actively proliferating cell population, including “*cell division”* and “*DNA replication initiation”* and contained >95% of cells isolated from muscles at 3 dpi (Figure 2C). All three clusters were enriched for genes linked to cell-cycle progression including *Cdk1* and *Ccne2* (Figure 2D and Table S2). Interestingly, cluster 0 exhibited a specific enrichment of several metabolic terms, including “*mitochondrial translation*” and “*tricarboxylic acid cycle*”. These results suggest that MuSCs activated *in vivo* contain a heterogenous population of proliferating cells. Dell’Orso and colleagues investigated the dynamic changes between quiescent MuSCs, *in vivo* activated MuSCs and proliferating myoblasts (PM) leveraging scRNAseq to provide a granular metabolic trajectory from quiescence to proliferation to differentiation (Dell’Orso et al., 2019). Metabolic pathways in general were found to be increased in both *in vivo* activated and PMs compared to quiescent MuSCs. Importantly, while a progressive enrichment in all metabolic pathways was observed, the dynamics and connectivity of genes in the individual metabolic pathways served as an identifier for individual cell states, supporting the notion that the metabolic blueprint of cells serves as a reliable differentiator and differences in metabolic profile are an indication of cellular heterogeneity.

Cluster 4 was enriched for terms linked to myogenic differentiation, including “*muscle contraction*” and “*skeletal muscle tissue regeneration*”, and had a significant enrichment of the myogenic differentiation factor *Myog* (Figure 2C,D and Table S2). It was interesting to note that like cluster 0, cluster 4 was similarly enriched for several metabolic terms including “*glycolytic process*” and “*oxidation-reduction process*” (Figure 2C). Together, these results suggest that the changing metabolic microenvironment experienced by MuSCs during skeletal muscle injury and repair is linked to significant changes in the innate metabolism of MuSCs. Importantly, despite all cells existing within the same local environment we observed significant differences in innate metabolism, linked to lineage progression and myogenic commitment.

Finally, cluster 2 contained a small population of cells that appeared to have exited the cell cycle and reached a differentiation checkpoint. Having failed this checkpoint, the cells have been marked for destruction, as evident from the enrichment of terms linked to apoptosis, including “*apoptotic process*” and “*release of cytochrome c from mitochondria*”.

To further investigate the link between innate cell metabolism and the process of myogenic commitment, we used immunofluorescence to label proliferating myogenic C2C12 cells with Pax7, MyoD (the master myogenic determination factor) and MitoTracker Red CMXRos (a marker of the proton gradient across the mitochondrial membrane, Figure S2A,B). Similar to Pax7^+^ MuSCs (Rocheteau et al., 2012), Pax7^+^ C2C12 cells exhibited a reduced level of MitoTracker red uptake (compared to Pax7^-^ cells, Figure S2C). In contrast, MyoD^+^ cells exhibited a high level of MitoTracker red uptake (Figure S2D). These results suggest a positive correlation between myogenic commitment and innate cell metabolism.

### Innate MuSC metabolism is dependent upon monosaccharide availability in the local microenvironment

To better understand the metabolic capabilities of proliferating myogenic cells we used a commercially available substrate phenotype array on C2C12 cells to compare monosaccharide utilization. We observed proliferating C2C12 cells utilize monosaccharides such as glucose and fructose with high efficiency, and with a significantly reduced capacity to process galactose and sorbose (Figure S3A). By using either high or low glucose concentrations, or replacing glucose with galactose, we were able to develop *in vitro* conditions of substrate limitation (low glucose) or processing limitation (galactose) to directly examine the role of the extracellular environment on innate MuSC metabolism.

Following 96 hrs of culture in glucose free DMEM supplemented with 25 mM glucose (high glucose, HG), 5 mM glucose (low glucose, LG) or 10 mM galactose (GAL), we measured the metabolic profile of MuSCs by performing a mitochondrial stress test on a Seahorse Bioanalyzer (Figure 3A). MuSCs cultured in LG exhibited a significant increase in the maximal oxygen consumption rate (OCR) compared to HG cultured cells (Figure 3B), indicating an increased oxidative capacity in these cells. These results are similar to those observed in MuSCs isolated from mice that underwent caloric restriction (Cerletti et al., 2012). In contrast to LG, MuSCs cultured in GAL exhibited a significant reduction in both basal and maximal oxygen consumption when compared to cells cultured in glucose. The decrease in basal OCR observed in MuSCs cultured in GAL was a result of decreased ATP synthase activity (Figure 3B). Concomitant measurements of the extracellular acidification rate (ECAR) revealed that while basal ECAR was not different between the three groups (Figure 3C,D), maximal ECAR was decreased significantly in MuSCs cultured in GAL compared to cells cultured in glucose (Figure 3D). Together, these changes in OCR and ECAR led to an increased metabolic capacity in LG cultured MuSCs and a decreased metabolic capacity in GAL cultured MuSCs (Figure 3E).

**Figure 3.**
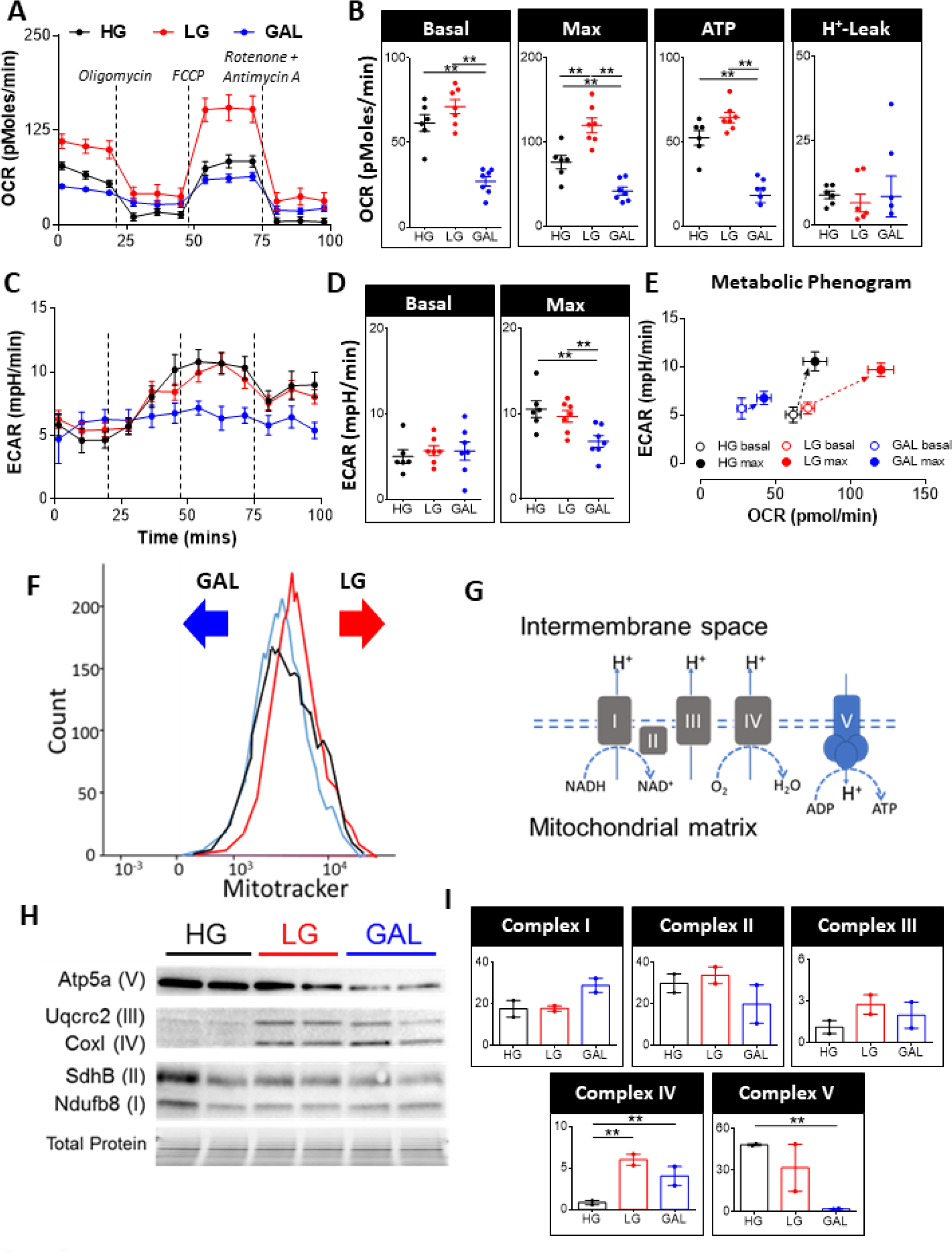
Innate MuSC Metabolism is Regulated via Extracellular Monosaccharide Availability. (A) A mitochondrial stress test was performed on MuSCs cultured for 96 hrs in high glucose (HG), low glucose (LG) or galactose (GAL) based growth media (n=6-7/group, mean ± SEM). (B) Quantification of basal and maximal respiration, and the rate of oxygen consumption used to generate ATP and H^+^-leak (determined from (A)). (C) The extracellular acidification rate (ECAR) as measured during (A). (D) Quantification of basal and maximal ECAR [determined from (C)]. (E) A metabolic phenogram demonstrating the altered metabolic capacity of cells cultured in HG, LG, or GAL. The length and direction of the arrow indicates maximal metabolic capacity. (F) Flow cytometric analysis of mitochondrial density in MuSCs cultured in HG, LG, or GAL (representative FACS plot presented, n=3 experiments performed). (G) Schematic outlining the role of each of the five mitochondrial complexes in the electron transport chain (ETC). (H) Western blot analysis of individual proteins within each of the five ETC complexes. (I) Quantification of (H) (n=2/group, mean ± SEM). ANOVA with Fisher’s LSD multiple comparison procedure, **p<0.01

Flow cytometric analysis of the mitochondrial proton gradient with MitoTracker Red CMXRos revealed a decrease in the proportion of high gradient MuSCs when cultured in GAL and an increase when cultured in LG (Figure 3F), supporting the observed changes in maximal oxidative capacity (measured on the Seahorse Bioanalyzer). We next examined the effects of altered monosaccharide availability on the expression of electron transport chain (ETC) proteins via western immunoblotting (Figure 3G,H). In support of a reduced capacity for the synthesis of ATP, MuSCs cultured in GAL exhibited a significant reduction in the Complex V protein ATP synthase subunit α compared to both HG and LG (Figure 3H,I). Interestingly, in contrast to the reduction in Complex V we observed an increased expression of Complex IV in MuSCs cultured in GAL (Figure 3H,I). These results also indicated a relative resistance for Complex I and II, which link to the TCA cycle, to undergo remodeling, suggesting an essential role for these complexes beyond electron transport and the generation of ATP.

### Monosaccharide availability within the extracellular environment regulates intracellular metabolite balance to control proliferation and global histone acetylation

As our data suggested a direct link between monosaccharide availability and the metabolic capacity of MuSCs, we next sought to investigate the intracellular metabolite balance of myogenic cells. To complete this analysis, we switched to an immortalized myogenic cell line (C2C12), which we confirmed underwent similar metabolic remodeling to MuSCs following culture in either HG, LG or GAL for 24 hrs (Figure S3B,C). Significant changes in the concentration of key glycolytic and TCA cycle metabolites were observed within 24 hrs of culture in either HG, LG or GAL (Figure 4A,B). Interestingly, cells cultured in GAL exhibited an elevated concentration of phosphoenolpyruvate (PEP), with a concomitant decrease in pyruvate. An accumulation of upstream glycolytic intermediates is typically linked to an increase in cell proliferation and is mediated by an isoform switch in pyruvate kinase from Pkm1 to the less active Pkm2 (Lunt et al., 2015; Ryall et al., 2015b). We observed that culture in GAL led to a reduction in the proportion of cells in the G1-phase of the cell cycle, with a concomitant increase in G2/M, leading to an overall increase in the rate of cell proliferation compared to HG cells (Figure S3D-F). This increase in proliferation was linked to a two-fold increase in the expression of the *pkm2* splice isoform which likely explains the observed build-up of PEP (Figure S3G). While cells cultured in LG did not exhibit a similar build-up of upstream glycolytic intermediates to GAL cells, we observed an increase in the proportion of cells in the S-phase of the cell cycle and an overall increase in cell proliferation (Figure S3D-F). This increase in cell proliferation in LG was associated with a decline in the expression of the active *pkm1* isoform (Figure S3G), indicating that cells cultured in either LG or GAL prioritize cell proliferation, with GAL cultured cells doing so at the expense of supplying the TCA cycle.

**Figure 4.**
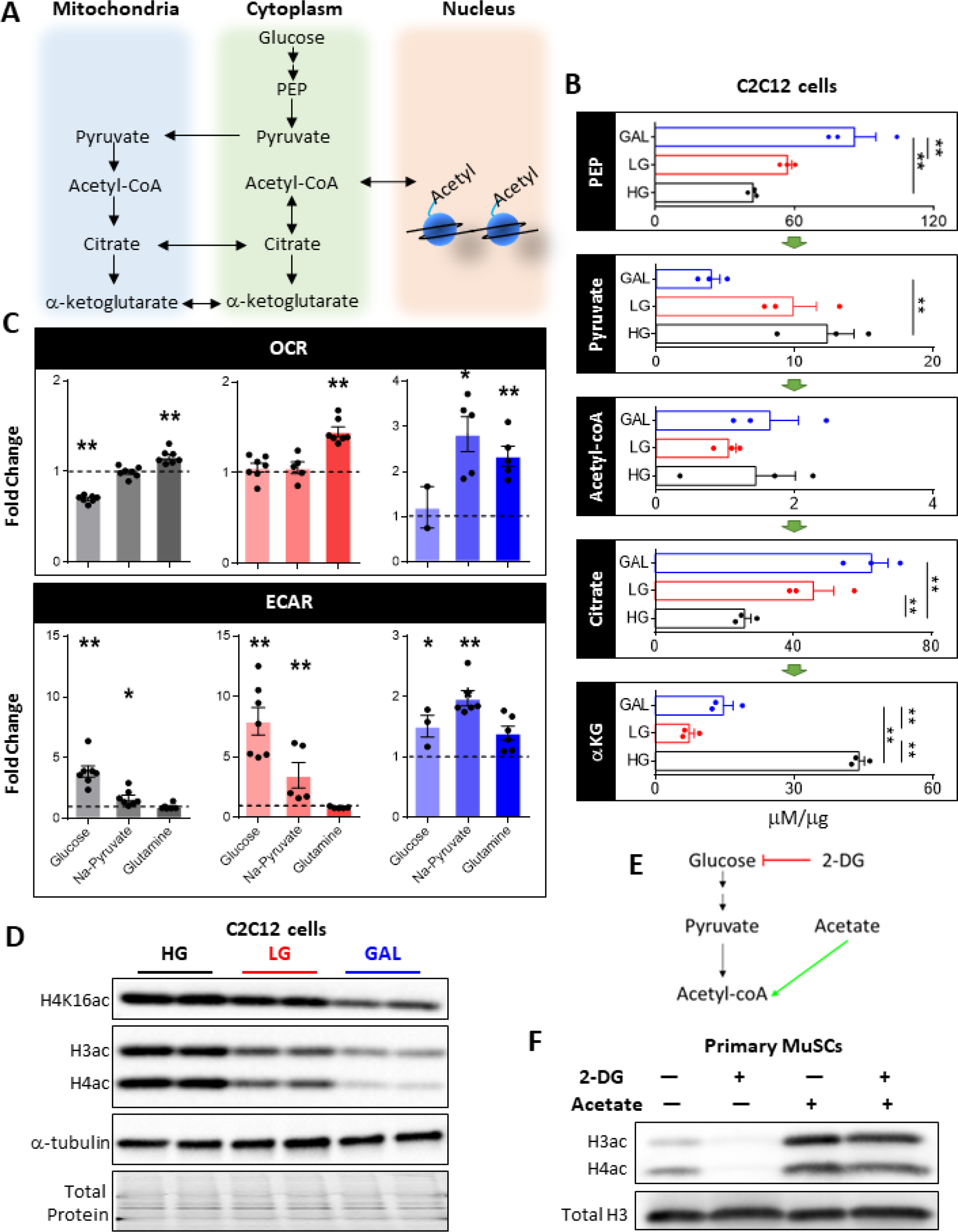
Monosaccharide Availability Regulates Cell Proliferation and Provides Temporal Regulation of Histone Acetylation in Myogenic Cells. (A) Schematic showing the important interplay between glycolysis, TCA cataplerosis and anaplerosis, and global histone acetylation. (B) Key glycolytic and TCA cycle metabolites were assessed in C2C12 cells utilizing colorimetric and fluorometric assays (n=3/group, mean ± SEM). (C) A Seahorse substrate utilization assay was performed on proliferating C2C12 cells cultured in either HG, LG, or GAL to determine the effect of substrate limitation on OCR and ECAR (n=6/group, data are presented as fold-change over substrate free media). (D) Western immunoblotting of specific (H4K16) and global histone acetylation in C2C12 cells (representative blot shown, n=4/group). (E) Schematic of 2-DG and acetate mechanism of action on histone acetylation. (F) Global histone acetylation in primary MuSCs in the presence or absence of the glycolytic inhibitor 2-DG and/or acetate (representative blot shown, n=4/group). ANOVA with Fisher’s LSD multiple comparison procedure, *p<0.05, **p<0.01

Despite a reduced concentration of pyruvate in cells cultured in GAL, we observed no significant difference in the concentration of acetyl-CoA among cells cultured in HG, LG or GAL and an increase in the concentration of citrate in cells cultured in LG or GAL (Figure 4B). To determine whether (and which) metabolites were rate-limiting, we performed a metabolite utilization assay on the Seahorse Bioanalyzer. Briefly, following 24 hrs of culture in either HG, LG, or GAL, we measured basal OCR and ECAR in C2C12 cells in media lacking glucose, sodium pyruvate and glutamine, and then measured the response after addition of either glucose, sodium pyruvate, or glutamine. All cells, regardless of preculture conditions, responded to the addition of glucose by switching to a glycolytic program, although this response was depressed in GAL cultured cells (Figure 4C). The addition of sodium pyruvate led to a small increase in ECAR in cells precultured in glucose, likely indicating that in these conditions the additional sodium pyruvate is converted to lactate. In contrast, the addition of sodium pyruvate to cells precultured in GAL led to a 2-3-fold increase in OCR suggesting that in these cells, sodium pyruvate is indeed rate-limiting for oxidative activity. Finally, glutamine led to a small but significant increase in OCR in cells precultured in glucose, but a robust 2-3- fold increase in cells precultured in GAL. When combined with our measures of intracellular metabolites, these data suggest a reduced capacity for cells precultured in GAL to oxidize glucose, and instead a reliance on glutamine to drive the TCA cycle and oxidative phosphorylation. Importantly, our data also demonstrate that while monosaccharide availability is limiting in LG cultured cells, these cells actually increase their overall metabolic capacity.

As we observed no decline in acetyl-CoA, despite a decrease in pyruvate, we hypothesized an additional carbon source for anaplerosis. Since one of the major (non-mitochondrial) acetyl-CoA stores is that linked to protein acetylation (Figure 4A), we hypothesized that cells cultured in GAL would maintain cell proliferation at the expense of protein acetylation (Zhao et al., 2016). To examine the possibility of acetylated proteins functioning as an energy store we analyzed global acetylation of histones H3 and H4 following culture in the three metabolic remodeling media (Figure 4D). As hypothesized, global acetylation of histones H3 and H4 was reduced significantly in GAL compared to HG cultured cells, suggesting an increase in citrate anaplerosis at the expense of histone acetylation. Interestingly, despite having an elevated metabolic capacity, we observed cells cultured in LG also exhibited a reduced level of histone acetylation. Similarly, we found H4K16ac (previously found to be enriched in activated MuSCs, Ryall et al., 2015b) was reduced in GAL, and to a lesser extent in cells cultured in LG compared to HG cultured cells (Figure 4D). While histone deacetylation in myogenic cells is regulated in a metabolic-dependent manner (Ryall et al., 2015b), here we demonstrate that global histone acetylation is directly regulated by extracellular carbohydrate availability rather than innate metabolic capacity.

To confirm a direct link between glycolysis and histone acetylation in proliferating myogenic cells we cultured primary MuSCs in HG alone or in the presence of the glucose transporter inhibitor 2-deoxy glucose (2-DG) or acetate, which can be converted into acetyl-CoA in a pyruvate-independent manner (Figure 4E). Inhibition of glucose uptake led to a dramatic decrease in the acetylation of H3 and H4 (Figure 4F). In contrast, the addition of acetate led to a significant increase in histone acetylation beyond basal levels. Importantly, administration of acetate circumvented the actions of 2-DG leading to a significant increase in histone acetylation (Figure 4F). These results support a role for extracellular carbohydrate availability in providing temporal regulation of global histone acetylation and chromatin permissiveness in dividing myogenic cells, and are consistent with findings showing that quiescent and differentiating MuSCs increase glucose utilization for respiration with consequently reduced acetylation (Yucel et al., 2019). In the study by Yucel and colleagues, proliferating MuSCs were enriched for the expression of H3K9ac, H4K16ac and H2BK5ac when compared to either quiescent or differentiating cells, allowing for the expression of genes associated with chromatin regulation, cell cycle regulation, and muscle cell proliferation. Proliferating MuSCs favored the incomplete oxidation of glucose to allow for the generation of acetyl-CoA, and in so doing served as a regulator of the transition between cells states through epigenetic landscape remodeling.

### Commitment to the myogenic lineage is regulated by monosaccharide availability in the local microenvironment

To determine the effect of monosaccharide availability on MuSCs, we first performed whole transcriptome sequencing (RNAseq) on freshly isolated MuSCs or on MuSCs that were cultured for 24, 48 or 72 hrs in growth media containing a high (25 mM) concentration of glucose. A principal component analysis revealed clear segregation in the transcriptional profiles of each timepoint (Figure 5A), while gene ontology analyses demonstrated a time-dependent progression through quiescence (*cell adhesion* and *negative regulation of cell proliferation* in freshly isolated MuSCs), activation (*translation initiation*, 24 hrs), proliferation (*cell division*, 48 hrs) and differentiation and fusion (*skeletal muscle tissue development*, 72 hrs, Figure 5B).

**Figure 5.**
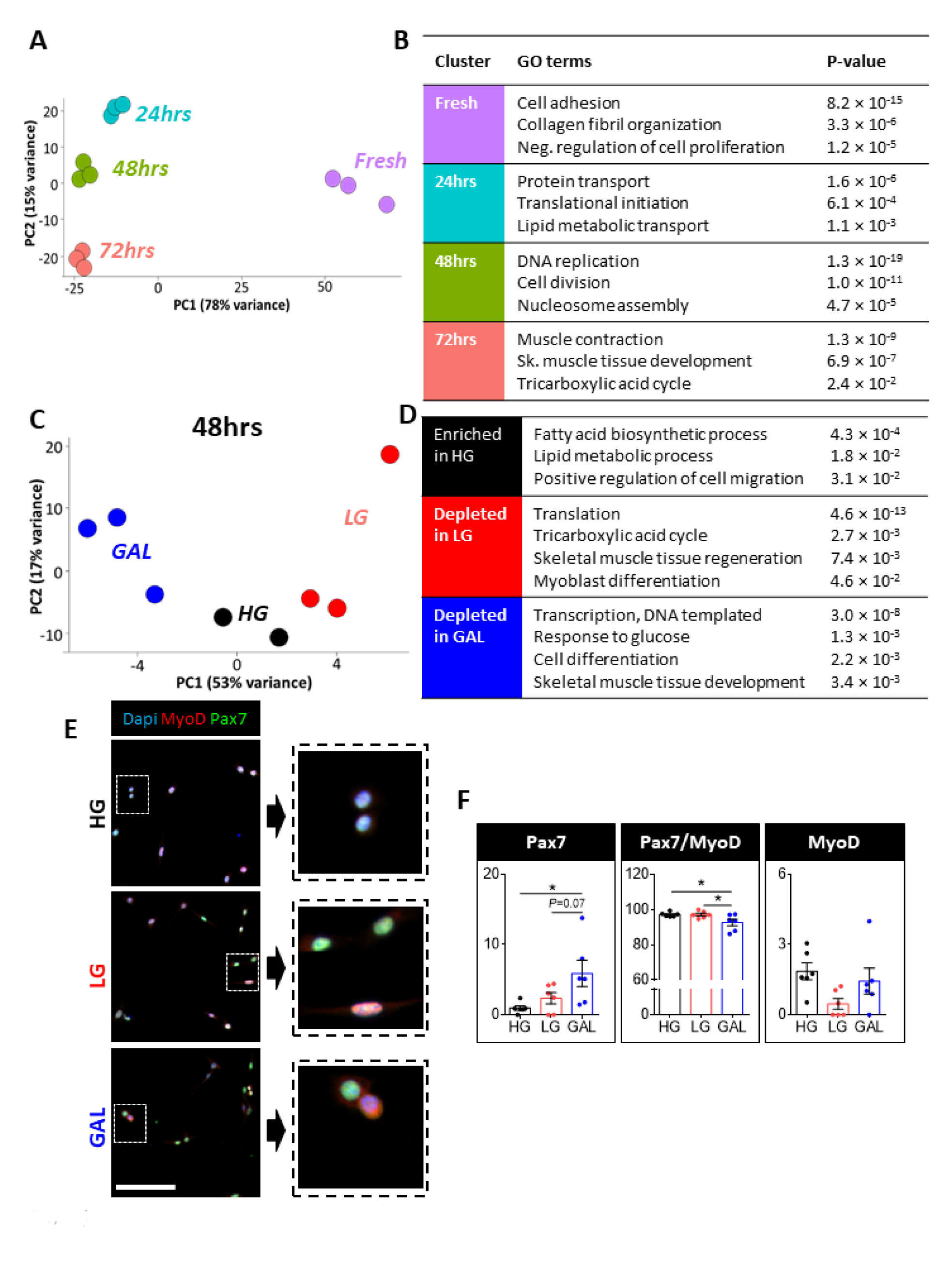
Commitment to the Myogenic Lineage Requires an Elevated Concentration of Extracellular Glucose. (A) Principal component analysis of gene expression performed on freshly isolated primary MuSCs, or MuSCs cultured in HG growth media for 24, 48 or 72 hrs (n=3/timepoint). (B) Results of gene-set enrichment analyses listing unique biological processes enriched at each timepoint. (C) Principal component analysis of gene expression performed on MuSCs following 48 hrs of culture in growth media containing HG, LG, or GAL (n=2-3/group). (D) Results of gene-set enrichment analyses for biological processes uniquely enriched (or depleted) in MuSCs from each group. (E) Immunofluorescence labelling of MuSCs cultured for 96 hrs in HG, LG, or GAL with Pax7 (green) and MyoD (red). (F) Quantification of (G) (n=6/group, mean ± SEM). ANOVA with Fisher’s LSD multiple comparison procedure, *p<0.05, **p<0.01

To better understand how changes in monosaccharide availability in the microenvironment might impact MuSCs, we cultured freshly isolated MuSCs in growth media supplemented with HG, LG or GAL for 48 hrs and then performed RNAseq. A principal component analysis revealed separation of the three groups (Figure 5C). Interestingly, MuSCs cultured in the presence of HG exhibited a specific enrichment of biological processes related to lipid metabolism (*fatty acid biosynthetic process* and *lipid metabolic process*) and the *positive regulation of cell migration* (Figure 5D). When the concentration of glucose was reduced or replaced with galactose, we observed a depletion of terms related myogenic progression, such as *skeletal muscle tissue regeneration* and *myoblast differentiation* (depleted in LG), and *cell differentiation* and *skeletal muscle tissue development* (depleted in GAL, Figure 5D).

To better examine the role of monosaccharide availability in the microenvironment on myogenic specification *per se*, we performed immunofluorescence labeling of MuSCs cultured for 96 hrs in HG, LG or GAL and characterized the proportion of MuSCs (Pax7^+^/MyoD^-^), activated MuSCs (Pax7^+^/MyoD^+^) and committed muscle progenitors (Pax7-/MyoD+, Olguin et al., 2007, Figure 4G). We found the proportion of Pax7^+^/MyoD^-^ MuSCs was elevated significantly in MuSCs cultured in GAL compared to HG (Figure 5F), confirming myogenic specification is directly regulated by monosaccharide availability and glycolysis.

### Single cell sequencing reveals the appearance of MuSC sub-populations that are influenced by the metabolic microenvironment

Our immunofluorescence analyses suggested population heterogeneity within each of the HG, LG, and GAL cultured groups. To resolve this heterogeneity, we again used our modified CELseq2 approach to perform single cell transcriptomics on over 450 FACS isolated MuSCs, either freshly isolated or following 96 hrs of culture in HG, LG or GAL conditions. In support of a robust transcriptional response to a change in the metabolic microenvironment, we observed that >50% of the top 100 most differentially expressed genes in HG, LG, and GAL cultured MuSCs were identically regulated in C2C12 cells (Figure 6A-C).

**Figure 6.**
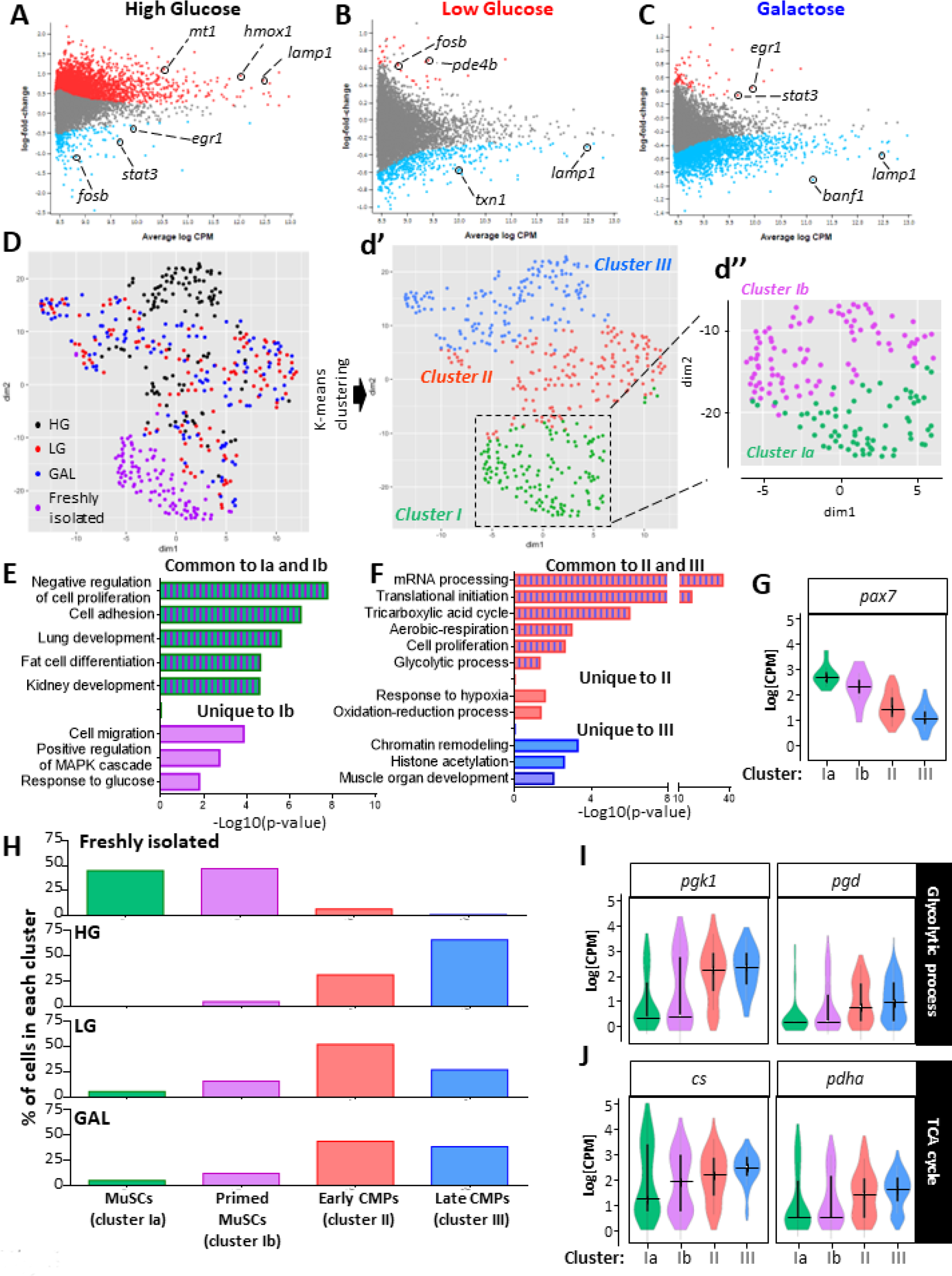
scRNA Sequencing Reveals Distinct Subpopulations of MuSCs which are Regulated by Extracellular Monosaccharide Availability. (A-C) MD plots examining the specific enrichment (red) or depletion (blue) of genes in MuSCs cultured for 96 hrs in HG (A), LG (B), or GAL (C), with a false discovery rate (FDR) cut-off of 0.05. (D) Single cells were mapped to two-dimensions using a t-SNE approach and three clusters were identified via a K-means clustering analysis (d’). Cluster I could further be subdivided into two clusters, Ia and Ib (d’’). (n=102-125 cells/group). (E-F) Common and unique biological processes enriched in clusters Ia and/or Ib (E), and clusters II and/or III (F). (G) Violin plot of detectable *pax7* expression in single cells. Horizontal bar indicates the median value and vertical bars extend from the 25^th^ to 75^th^ percentiles. (H) Proportion of freshly isolated MuSCs, or cells cultured for 96 hrs in either HG, LG, or GAL identified within each of the four clusters. (I-J) Violin plots of gene expression for glycolytic genes (I) and TCA cycle enzymes (J). Horizontal bar indicates the median value and vertical bars extend from the 25^th^ to 75^th^ percentiles. ANOVA with Fisher’s LSD multiple comparison procedure, *p<0.05, **p<0.01

To visualize cell population dynamics, we again examined heterogeneity using tSNE with K-means clustering (Figure 6D). Utilizing this larger sample size, we identified three distinct clusters of cells (Figure 6d’), with cluster I further divided into two sub-clusters (Cluster Ia and Ib, Figure 6d’’). Cluster I cells exhibited an enrichment of genes associated with developmental terms, *cell adhesion* and *negative regulation of cell proliferation*, common to freshly isolated MuSCs. Interestingly, cluster Ib was specifically enriched for terms associated with a more activated or ‘poised’ state including *response to glucose*, *positive regulation of MAPK cascade* and *cell migration* (Figure 6E). In contrast to cluster I, clusters II and III were both enriched for metabolic terms typical of actively proliferating MuSCs, including *tricarboxylic acid cycle*, *aerobic respiration,* and *glycolytic process* (Figure 6F). While cluster III was also enriched for terms such as *histone acetylation* and *muscle organ development*, cluster II was specifically enriched for *chromatin silencing* and *telomere maintenance* (Figure 6F), suggesting a feedforward model whereby the microenvironment can directly regulate chromatin structure and expression of genes that, in turn, regulate chromatin structure. Importantly, an examination of *pax7* expression revealed that cells in clusters Ia and Ib exhibited the highest level of expression, and clusters II and III the lowest expression (Figure 6G). Based on both gene ontology and *pax7* expression, we classified each cluster as either MuSCs (cluster Ia), primed MuSCs (cluster Ib), early committed myogenic progenitors (CMPs, cluster II), or late CMPs (cluster III).

We next examined the contribution of freshly isolated and HG, LG and GAL cultured cells to each cluster and found that freshly isolated MuSCs were located predominantly in cluster I with an equal proportion in cluster Ia and Ib, and a minor contribution to cluster II (Figure 6H). Clusters Ia and Ib consistently exhibited lower expression of key metabolic enzymes (Figure 6I, J). The majority of HG cultured cells populated cluster III with a small proportion in cluster II, and less than 5% in cluster Ib. In contrast, the majority of cells cultured in LG and GAL were found in cluster II with ∼20% of LG and GAL cells also found in clusters I and III (Figure 6H). Interestingly, both LG and GAL groups contained a small population of cells (∼5%) in cluster Ia that was never observed in the HG group (Figure 6H). The study of single MuSC transcriptomes has revealed a previously unappreciated level of heterogeneity in both freshly isolated and proliferating MuSCs. Importantly, we have shown that the progression from MuSC through late CMP is regulated in a metabolic-dependent manner, with a “*response to glucose*” identified as an early marker of MuSC activation.

### The metabolic microenvironment regulates myogenic commitment and differentiation of MuSCs

To confirm our single cell immunofluorescence and scRNAseq results that were indicative of increased MuSC self-renewal, we next examined the effect of altered carbohydrate availability on myogenic specification in an *ex vivo* single fiber culture model, which allows for the study of MuSCs within their intact niche. Single myofibers isolated from the extensor digitorum longus (EDL) muscles of C57Bl6 mice were cultured in HG, LG, or GAL media for 40, 72, or 96 hrs (Figure 7A). When cultured in HG-based media, MuSCs rapidly induced MyoD expression with greater than 98% of cells expressing MyoD after 40 hrs of culture and less than 2% of cells retaining a MuSC (Pax7^+^/MyoD^-^) phenotype after 96 hrs of culture (Figure 7B). Fibers cultured in LG exhibited a reduced proportion of MyoD^+^ cells with approximately 4% of cells persisting in a MyoD^-^ MuSC state after 96 hrs of culture (Figure 7B). More remarkably, when cultured in GAL-based media, more than 20% of cells maintained a MyoD^-^ MuSC phenotype at 40 hrs of culture, and this population remained elevated at 72 and 96 hrs of culture (Figure 7B). Importantly, the increased proportion of Pax7^+^ cells on GAL cultured fibers was a result of an increase in the absolute number of Pax7^+^ cells (Figure 7C).

**Figure 7.**
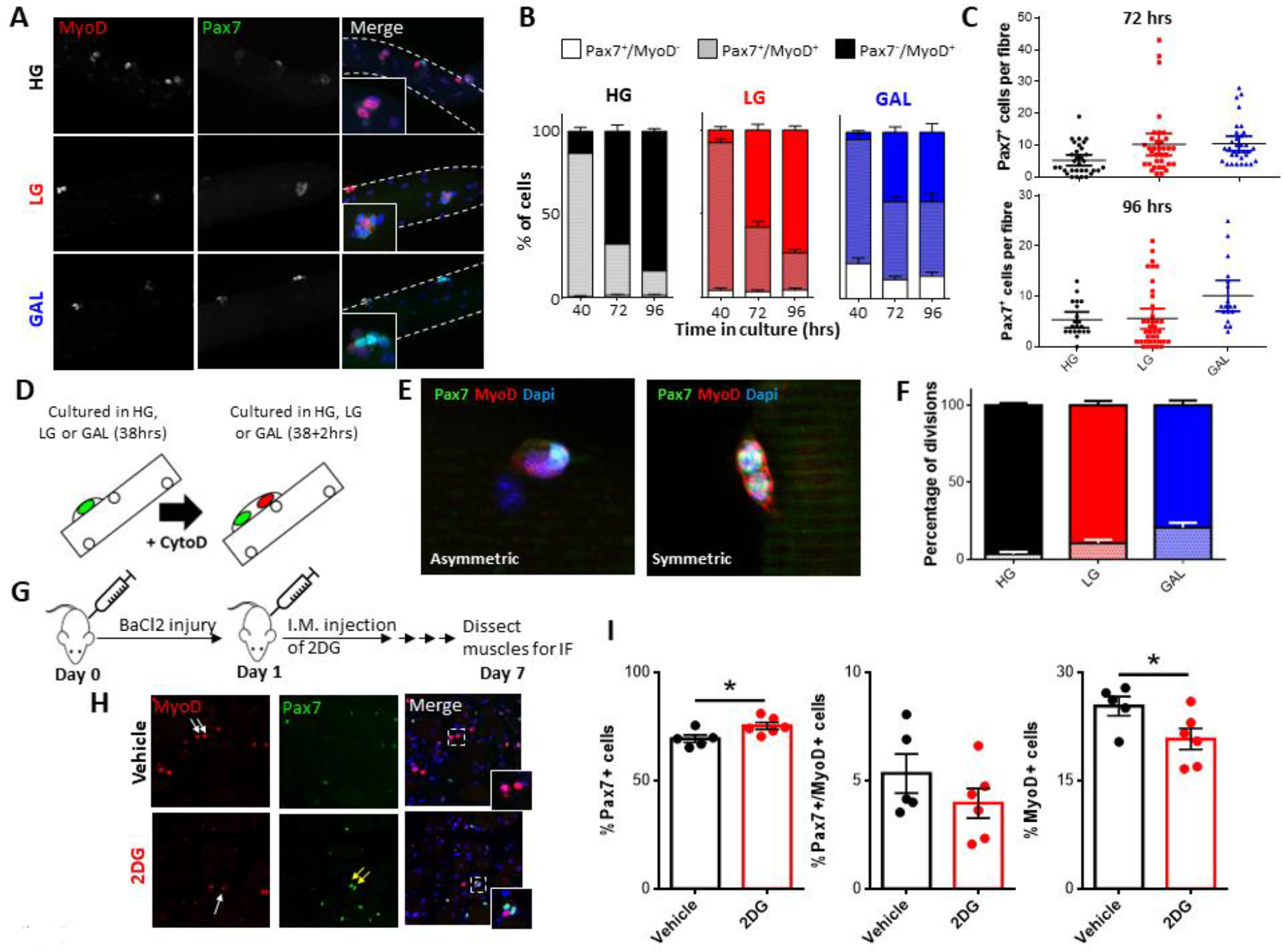
Monosaccharide Availability Regulates the Balance Between MuSC Self-Renewal and Myogenic Commitment. (A) Single muscle fibers were cultured in HG, LG, or GAL and immunofluorescently labelled with Pax7 and MyoD. (B) Quantification of the proportion of self-renewing MuSCs (Pax7^+^/MyoD^-^), activated MuSCs (Pax7^+^/MyoD^+^) and committed muscle progenitors (Pax7^-^/MyoD^+^) on fibers cultured for 40, 72, or 96 hrs in HG, LG or GAL (n>80 fibers/timepoint, isolated from n=6 mice, mean ± SEM). (C) Quantification of the absolute number of Pax7^+^/MyoD^-^ MuSCs in (B) at 72 hrs and 96 hrs. (D) Schematic detailing the use of cytochalasin D to directly analyze MuSC asymmetric division. (E) Representative images of an asymmetrically-and a symmetrically-dividing MuSC following two-hours of culture in cytochalasin D and immunofluorescently labeled with Pax7 and MyoD. (F) Quantification of (E). (n>50 fibers, isolated from n=2 mice, mean ± SEM). (G) Schematic of the injury protocol with (or without) subsequent inhibition of glucose uptake. (H) Representative images of regenerating TA muscles seven days post-injury. White arrows indicate Pax7-/MyoD+ muscle progenitors, and yellow arrows indicate Pax^+^/MyoD^-^ MuSCs. (I-K) Quantification of the percent of (I) Pax7^+^ cells, (J) Pax7^+^/MyoD^+^ cells, and (K) MyoD^+^ cells (n=5-6/group).

In addition to regulating myogenic commitment, we found that extracellular carbohydrate availability could regulate differentiation (Figure S4A). Single myofibers cultured in GAL-based growth media exhibited a decrease in the proportion of committed progenitors that entered the differentiation program (determined via the expression of myogenin) when compared to myofibers cultured in HG or LG conditions (Figure S4B), again supporting an essential role for the metabolic microenvironment in myogenic lineage progression.

### The metabolic microenvironment promotes MuSC self-renewal via increased asymmetric division

To better understand the mechanism through which the metabolic microenvironment could maintain an increased proportion of Pax7^+^ MuSCs, we therefore examined the types of divisions MuSCs underwent, focusing on characterizing the proportion of asymmetric and symmetric divisions. A Pax7^+^ MuSC can undergo one of three types of divisions: asymmetric stem cell maintenance; symmetric stem cell expansion; or symmetric progenitor expansion (Dumont et al., 2015). It has been proposed that asymmetric divisions maintain a healthy stem cell pool while unregulated symmetric stem cell expansion or progenitor expansion can result in pathological stem cell hyperplasia or depletion of the MuSC pool, respectively (Dumont et al., 2015). We cultured single fibers for 40 hrs to allow the majority of MuSCs to undergo first division and examined the differential expression of Pax7 and MyoD in daughter cells (Figure S4C). In support of our analysis of Pax7/MyoD over 96 hrs, when cultured in GAL-based growth media the proportion of asymmetric divisions was increased more than two-fold compared to single fibers cultured in LG and HG (Figure S4D,E), as determined by the differential expression of Pax7 and MyoD in daughter cells.

As MuSCs are a highly mobile population of cells and move rapidly along the fiber to which they are attached when cultured (Siegel et al., 2009), we sought to more robustly characterize asymmetric division using cytochalasin D (CytoD) an inhibitor of cytokinesis. Culturing proliferating cells in the presence of CytoD allows for nuclear but not cell division (Figure 7D), facilitating confirmation of asymmetric division (Casella et al., 1981). Live cell imaging of single fibers cultured in the presence of CytoD also confirmed that CytoD inhibited movement of MuSCs (data not shown). Single fibers were cultured in HG-, LG-, and GAL-based media for 38 hrs (prior to first division) before being replaced with media containing 1 mM CytoD (Figure 7D). This clearly identified both symmetric and asymmetric divisions of MuSCs (Figure 7E) and revealed a two-and five-fold increase in the rate of asymmetric divisions observed in fibers cultured in LG and GAL, compared to HG (Figure 7F).

To determine whether the local elevation of glucose at the site of muscle injury is directly linked to the commitment of MuSCs to the myogenic lineage, we returned to employing the BaCl_2_ muscle injury model in mice. Twenty-four hours after BaCl_2_-induced injury we performed an intramuscular injection of 2DG (or vehicle) and then excised the muscles 7-d later (Figure 7G). By this time, the majority of MuSCs have undergone multiple rounds of proliferation and stopped expressing MyoD before their return to quiescence (Figure 7H). We observed a significant increase in the proportion of Pax7^+^/MyoD^-^ MuSCs in muscles that had been treated with 2DG, with a concomitant decrease in the proportion of committed myoblasts (Figure 7I-K). These results provide the first clear evidence for monosaccharide availability in the microenvironment regulating MuSC self-renewal and support a direct link between elevated glucose availability and committed progenitor expansion.

## DISCUSSION

In this study we have shown that extracellular monosaccharide availability is a key source of carbon for histone acetylation and myogenic lineage progression. Importantly, we have shown that the master myogenic regulator MyoD is exquisitely sensitive to changes in glucose uptake and pyruvate oxidation. Under substrate limited conditions, MuSCs prioritize cell division over histone acetylation, leading to an increased rate of asymmetric cell division and self-renewal. These results support a key role for the metabolic microenvironment in the temporal regulation of histone acetylation and myogenic commitment.

Activation of MuSCs has been linked to increased glycolysis, a decrease in [NAD^+^], and a subsequent decrease in the histone deacetylase activity of SIRT1 (Ryall et al., 2015b). This decrease in SIRT1 activity was found to promote acetylation of H4K16 and increase transcription of *Myod1*. Here we show that in addition to reducing SIRT1 deacetylase activity, increased glycolysis can provide the necessary carbon for histone acetylation. Importantly, our results support a model whereby extracellular monosaccharide availability rather than innate metabolic capacity regulates histone acetylation, as cells cultured in LG conditions exhibited an increased metabolic capacity but reduced histone acetylation. These results provide strong evidence that the extracellular environment can provide temporal regulation to histone acetylation and transcription.

The first link between metabolism and lineage progression in MuSCs was identified by Rocheteau and colleagues (2012) who found that Pax7^Hi^ cells displayed reduced intensity of labelling for MitoTracker compared to Pax7^Lo^ cells. This inverse link between mitochondrial content and MuSC “stemness” was strengthened further by several studies demonstrating a gradual increase in mitochondrial content as MuSCs shift from quiescence to an activated state (Rodgers et al., 2014; Ryall et al., 2015b). Using scRNAseq we have shown that freshly isolated *Pax7*^Hi^ MuSCs display a homologous low metabolic phenotype. In response to *in vitro* activation MuSCs become enriched for key glycolytic, TCA, and oxphos regulators, and exhibit reduced levels of *Pax7*. However, a subpopulation of activated MuSCs maintain (or revert to) a low metabolic phenotype and are phenotypically similar to freshly isolated MuSCs. Heterogeneity within transcription factors (Myf5, Myod1, Pax7, others), RTK inhibitors (Sprouty1) and structural proteins (Dmd1) in proliferating MuSCs has been well described (Cheung et al., 2012; Dumont et al., 2015; Kuang et al., 2007; Shea et al., 2010). Our results suggest that innate cell metabolism is also heterogeneous, with highly metabolic MuSCs more likely to undergo commitment to the myogenic lineage.

It was suggested previously that glycolysis might regulate MuSC self-renewal and return to quiescence (Theret et al., 2017), but we have shown here that MyoD is exquisitely sensitive to the availability of glucose and histone acetylation. Inhibition of glucose uptake via culture with 2-DG led to a dose-dependent decrease in both global histone acetylation and MyoD protein levels. Importantly, Theret and colleagues found a link between decreased oxidative capacity and MuSC self-renewal (Theret et al., 2017). As we have identified a central role for the TCA cycle in the regulation of histone acetylation and myogenic commitment, it is essential that future studies focus on metabolism and MuSC self-renewal include an analysis of TCA metabolites such as acetyl-CoA and citrate.

By combining a large number of single cell transcriptomes from freshly isolated and cultured MuSCs we identified four novel populations of cells, two which were composed predominantly of freshly isolated cells, and two composed of cultured cells. In the two populations composed primarily of freshly isolated cells, we observed an enrichment of GO terms such as cell migration and response to glucose which are similarly enriched in MuSCs that have been activated for 5 hrs (compared to *in vivo* fixed cells believed to be truly quiescent MuSCs; Machado et al., 2017). It has become increasingly recognized that quiescence is a heterogeneous state with quiescent MuSCs capable of adaptively moving to a shallower state of quiescence following injury at a distant site (G_Alert,_ Rodgers et al., 2015). Therefore, we propose that quiescent MuSCs contain a heterogenous population of cells, with those expressing the highest levels of Pax7 less likely to undergo a shift to the primed state, a state of potential “deep quiescence”. This model is similar to that proposed by Rocheteau and colleagues who observed that Pax7^Hi^ cells were a metabolically low population that exhibited a prolonged time to first division (Rocheteau et al., 2012).

Similar to freshly isolated MuSCs, cultured MuSCs primarily separated into two clusters termed “*early*” and “*late*” CMPs. While both clusters exhibited an enrichment of metabolic terms when compared to cluster I, only the *late CMP* cluster was enriched for muscle differentiation terms. Importantly, we did not detect the late differentiation marker, myogenin, in any cell, suggesting that the *late CMP* cluster marked a population of cells that had not yet undergone differentiation. Interestingly, we found that monosaccharide availability controlled the proportion of MuSCs in the *early* and *late CMP* clusters, a result that was directly supported by our single fiber culture model that confirmed MuSCs cultured in either LG or GAL exhibited a reduced (or delayed) capacity to undergo specification and differentiation.

Muscle regeneration is a highly complex and precisely coordinated process. Detailed characterizations have been made about the role of neighboring cells, inflammation, cytokines, and growth factors in regulating this process, but the role of metabolites has only recently gained attention. For example, endothelial-derived lactate has been flagged for its crucial role in polarizing M2 like macrophages to support angiogenesis at the local injury site to allow for muscle regeneration (Zhang et al., 2020). Macrophages themselves excrete glutamine into the local interstitial space that then is taken up by the SLC1A transporter into MuSCs, initiating mTOR signaling and enhancing muscle regeneration and self-renewal of MuSCs (Shang et al., 2020). In the present study, we show that muscle injury is accompanied by localized hyperglycemia and that blockade of monosaccharide utilization by 2DG results in an increase in Pax7^+^/MyoD^-^ MuSCs and a decrease in committed progenitors. Our results provide a model whereby localized tissue hyperglycemia after injury regulates transcription and myogenic lineage progression. Together, these studies highlight a crucial role for metabolites in the regulation of muscle regeneration and MuSC lineage progression.

The data presented here document an important role for the extracellular metabolic environment in the regulation of MuSC lineage progression and cell fate. Specifically, we have demonstrated that extracellular glucose is necessary to support histone acetylation and commitment to the myogenic lineage. In the absence of a high concentration of glucose, MuSCs prioritize cell division over histone acetylation, leading to asymmetric cell division and self-renewal. In this model, we have identified pyruvate oxidation and the TCA cycle as key regulators of the balance between stem cell self-renewal and progenitor expansion.

## STAR METHODS

## KEY RESOURCES TABLE

## CONTACT FOR REAGENT AND RESOURCE SHARING

Further information and requests for resources and reagents should be directed to and will be fulfilled by the Lead Contact, James Ryall

## EXPERIMENTAL MODEL AND SUBJECT DETAILS

### Mice

All procedures were approved by the Animal Ethics Committee of The University of Melbourne and conformed to the Australian code of practice for the care and use of animals for scientific purposes as stipulated by the National Health and Medical Research Council of Australia. Mice were housed in the Biological Research Facility at The University of Melbourne under a 12h light-dark cycle, with drinking water and standard chow provided *ad libitum*. The Pax7^creERT2^R26-eYFP^fl/fl^ mice were generated from Pax7^creERT2^ (Murphy et al., 2011) and R26-eYFP^fl/fl^ (Srinivas et al., 2001) founder mice strains.

### C2C12 cells

C2C12 cells purchased from the ATCC were cultured at 37°C in a normoxic environment maintained with 5% CO_2_. Growth media contained Dulbecco’s Modified Eagle Medium (DMEM, cat# 11966025, Gibco) supplemented with 1 mM sodium pyruvate (cat# 11360070, Gibco), 10% fetal bovine serum (FBS, Australian origin, cat# 10099141, Gibco), 1% penicillin/streptomycin (Gibco) and either 25 mM or 5 mM glucose, or 10 mM galactose.

### Primary MuSC Culture

Primary MuSCs were cultured at 37°C in a normoxic environment maintained with 5% CO_2_. Growth media contained Dulbecco’s Modified Eagle Medium (DMEM, cat# 11966025, Gibco) supplemented with 1 mM sodium pyruvate (cat# 11360070, Gibco), 10% fetal bovine serum (FBS, Australian origin, cat# 10099141, Gibco), 10% horse serum (HS, New Zealand origin, cat# 16050130, Gibco), 0.05% chick embryo extract (cat# C3999, US Biological), 2.5 ng/ml basic fibroblast growth factor (bFGF, cat# 100-18B, PeproTech), 1% penicillin/streptomycin (Gibco) and either 25 mM or 5 mM glucose, or 10 mM galactose.

## METHOD DETAILS

### Whole muscle metabolomics

To induce injury, mice received a 40μL intramuscular injection of 1.2% barium chloride in saline into the right TA muscle. Following induction of muscle injury, mice were placed in a recovery cage and carefully monitored until they had recovered from anaesthesia. On the day of day of dissection, mice were killed via rapid cervical dislocation, and the right TA muscle immediately harvested. The TA muscle was quickly quenched in liquid nitrogen to halt metabolism and stored at -80°C for later metabolomic analyses.

All the following steps were performed with pre-chilled solutions and completed in the presence of dry ice. A solution comprising methanol and water (80:20) containing internal standards (13C-AMP, 13C-UMP, 13C Sorbitol, and 13C Valine) was added to the snap-frozen TA muscle samples which were then mechanically homogenised using a micro homogenizer connected to a pre-chilled pestle. Samples were vortexed and sonicated for five minutes to ensure complete cell lysis and then incubated for five minutes at 4°C before separation into a polar metabolite fraction for steady-state analysis.

For measurement of steady-state polar metabolomics, samples were centrifuged (Eppendorf 5810 R, Hamburg Germany) at maximum speed for 10 minutes at 4°C prior to injection into a hydrophilic interaction (HILIC) liquid chromatography (LC) and high-resolution mass spectrometer (QTOF-6545, Agilent, Santa Clara, CA, USA). Metabolite identification and quality checks for polar metabolomics was performed using QTOF MassHunter Quant software (Agilent, Santa Clara, CA, USA).

### Matrix-assisted laser desorption/ionization (MALDI) mass spectrometry imaging (MSI)

Frozen tissues were cryosectioned at 12 µm thickness using a Leica CM1860 Cryomicrotome (Leica Microsystems Pty Ltd, Mt Waverly, VIC, Australia), then thaw mounted to glass slides. Samples were vacuum dried in a desiccator for a minimum of 15 min prior to matrix deposition using a TM-Sprayer (HTX Technologies, Carrboro, NC). The spray settings for norharmane (5 mg/mL in 98% acetone) with added sodium 2- anthraquinonesulfonate (500 ng/mL) application were: solvent flow rate 0.15 mL/min, nozzle speed 1200 mm/min, nozzle temperature 30°C, track spacing 2 mm, 8 passes with 90° rotation on passes 2, 4, 6, and 8, with 1 mm track offset for passes 3, 4, 7, and 8. Following matrix deposition, samples were stored in a vacuum desiccator until data collection.

A hybrid MALDI/ESI 7 Tesla XR SolariX MRMS (Bruker, Bremen, Germany) was used for data acquisition. The instrument was operated in negative ionisation mode using optimized conditions for mass transfer between *m/z* 70−2000 with a resolving power of 100,000 at *m/z*400. Data were acquired using FlexImaging 5.0 and ftmsControl 2.2 software (Bruker, Bremen, Germany). Laser settings included a minimum spot size, between 200 shots with laser power set to 35%, a raster of 50 × 50 µm, Smart Walk Enabled with a width of 45 µm. The instrument was calibrated with elemental red phosphorous clusters prior to acquisition. Lock mass for 2- anthraquinonsulfonate (*m/z* 287.001968) was used for mass calibration during acquisition.

Statistical analysis and visualization were performed in SCiLS Lab 2019c software (Bruker, Bremen, Germany). Discriminant analysis was conducted by calculating Receiver Operating Characteristic (ROC) curves and the Area Under the Curve (AUC) Data. Highly discriminant peaks were selected at thresholds of 0.1 and 0.9 AUC to reduce the total number of peaks for analysis. Data were RMS normalized with signal intensity auto-scaled and visualised using the Viridis color map. Datasets were exported from SCiLS Lab in the common imzML format and uploaded to Metaspace (www.metaspace2020.eu) for automated metabolite annotation. Individual accurate masses were calculated for specific metabolites and tentative annotations provided based on mass errors of less than 2 ppm.

In negative ionization mode, hexose sugar was observable as the deprotonated negative ion ([M-H)^-^ *m/z* 179.05615, 0.2 ppm mass error) and was also strongly discriminant for the injured tissue (AUC = 0.138). A phosphorylated hexose species was observed as the deprotonated ion ([M-H)^-^ *m/z* 259.02288, 1.7 ppm mass error) and further dehydrated hexose-phosphate was also observed with a similar spatial distribution to the hexose-phosphate ([M-H)^-^ *m/z* 241.01219, 1.3 ppm mass error). The hexose-6-phosphate was strongly discriminant for healthy tissues (AUC = 0.869), whereas the dehydrated analogue was not discriminative.

### MuSC Isolation

For the FACS isolation of eYFP^+^ MuSCs, 9-week-old male Pax7^creERT2^R26-eYFP^fl/fl^ mice were treated with tamoxifen (20 mg/ml in corn oil) via daily i.p. injections (100 μL) for five consecutive days. Two weeks after the final injection, animals were killed and the hindlimb skeletal musculature was collected and dissociated as described previously (Liu et al., 2015). Cells were sorted on a FACS Aria III (Becton Dickinson Biosciences, New Jersey, USA), gating on YFP positivity.

### CELseq2 Analysis of Single MuSCs

Single MuSCs were flow sorted into a chilled 384-well PCR plate (Greiner, 785290) containing 1.2 μL of primer/lysis mix [20 nM indexed polydT primer (custom made, IDT), 1:6,000,000 dilution of ERCC RNA spike-in mix (Ambion – 4456740), 1 mM dNTPs (NEB - N0446S), 1.2 units SUPERaseIN Rnase Inhibitor (Thermo Fisher – AM2696), DEPC water (Thermo Fisher – AM9920)] using a BD FACSAria III flow cytometer (BD Biosciences, San Jose, CA, USA). Sorted plates were sealed, centrifuged for 1 min at 3000 rpm and immediately frozen upside down at -80°C until further processing using an adapted CELSeq2 protocol (Hashimshony et al., 2016).

Sorted plates were thawed on ice and briefly centrifuged. To lyse the cells and anneal the mRNA capture primer, the plate was incubated at 65°C for 5 min and immediately chilled on ice for at least 2 min before adding 0.8 μL reverse transcription reaction mix [in 2 μL RT reaction: 1× First Strand buffer (Invitrogen – 18064-014), 20 mM DTT (Invitrogen – 18064- 014), 4 units RNaseOUT (Invitrogen 10777-019), 10 units SuperScript II (Invitrogen 18064- 014)]. The plate was incubated at 42°C for 1 hr, 70°C for 10 min and chilled to 4°C to generate first strand cDNA. For second strand cDNA synthesis 6 μL of second strand reaction mix was added [1× NEBNext Second Strand Synthesis buffer (NEB #E6111S), NEBNext Second Strand Synthesis Enzyme Mix: 2.4 units DNA Polymerase I (*E. coli*), 2 units RNase H, 10 units *E. coli* DNA Ligase (NEB #E6111S), DEPC water (Thermo Fisher – AM9920)]. The plate was incubated at 16°C for 2 hrs to generate double stranded cDNA.

All samples were pooled and cleaned using a 1.2× NucleoMag NGS clean-up and size select magnetic beads (Macherey-Nagel – 7449970.5) according to manufacturer’s instructions. To reduce the number of beads for each 100 μL pooled sample, 20 μL beads and 100 μL bead binding buffer (20% PEG8000, 2.5M NaCl, pH5.5) was added. The cDNA was eluted in 6.4 μL DEPC water and combined with 9.6 μL of IVT reaction mix [1.6 μL of each of the following: A,G,C,U, 10× T7 buffer, T7 enzyme (MEGAscript T7 Transcription Kit – Ambion AM1334)] and then incubated at 37°C for 13 hrs and then chilled and kept at 4°C.

To remove leftover primers 6 μL ExoSAP-IT For PCR Product Clean-Up (Affymetrix – 78200) was added and the sample was incubated at 37°C for 15 min and then chilled and kept at 4°C.

Chemical heat fragmentation was performed by adding 5.5 μl of 10× Fragmentation buffer (RNA fragmentation reagents – AM8740) to the sample which was then incubated in a pre-heated thermal cycler at 94°C for 2.5 min and then immediately chilled on ice. Following this, 2.75 μL of Fragmentation Stop buffer (RNA fragmentation reagents – AM8740) was added and the fragmented amplified RNA was purified using 1.8× RNAClean XP beads (Beckman coulter – A63987) according to manufacturer’s instructions and eluted in 6 μL DEPC water. 5 μL of fragmented RNA was transcribed into cDNA using 5’-tagged random hexamer primers (GCCTTGGCACCCGAGAATTCCANNNNNN) introducing a partial Illumina adapter as also described in CELseq2 (Hashimshony et al., 2016). To remove RNA secondary structure and anneal the mRNA capture primer 1 μL of tagged random hexamer (100 μM) and 0.5 μL of 10 mM dNTPs (dNTP solution set NEB - N0446S) were added to the sample, incubated at 65°C for 5 min and then immediately chilled on ice for at least 2 min before adding 4 μL reverse transcription reaction mix [in 10 μL RT reaction: 1× First Strand buffer (Invitrogen – 18064- 014), 20 mM DTT (Invitrogen - 18064-014), 4 units RNaseOUT (Invitrogen 10777-019) and 10 units SuperScript II (Invitrogen 18064-014)].

The PCR primers introduce the full-length adaptor sequence required for Illumina sequencing (for details see Illumina small RNA PCR primers). PCR was performed in 12.5 μL using half of the ranhexRT sample as a template [1× KAPA HiFi HotStart ReadyMix (KapaBiosystems KK2602), 400 nM each primer].

The final PCR amplified library was submitted to two consecutive 1× NucleoMag NGS Clean-up and Size select magnetic bead selections (Macherey-Nagel – 7449970.5) according to the manufacturer’s instructions. The final library was eluted in 20 μL of 10 mM Trizma hydrochloride solution (Sigma-Aldrich – T2319-1L) and 50 bp paired-end sequencing was performed on an Illumina NextSeq 550.

Read pairs from the CELSeq2 scRNA-sequencing were mapped to the mm10 mouse genome using the Subread aligner (Liao et al., 2013), and assigned to genes using scPipe with ENSEMBL v86 annotation. Gene counts were exported by scPipe with UMI-aware counting and imported into R (Tian et al., 2018). Samples were filtered using the detect_outlier function from scPipe and removed if they were detected as outliers or had fewer than 1,000 total genes detected. Genes that failed to achieve 1 count in at least 3 cells or 20 counts across all cells were filtered from the analyses. Batch effect across plates was removed using ComBat from the sva package (Johnson et al., 2007).

Dimensionality reduction was performed on normalized log2-cpm expression values with size factors from computeSumFactors in scran (Lun et al., 2016). t-SNE plots were generated with the Rtsne package using correlation distance between cells. Clustering was performed using k-means on the normalized expression values. Mean-difference (MD) plots were generated using limma (Ritchie et al., 2015) and Glimma (Su et al., 2017). Differential expression analyses used likelihood ratio tests with negative binomial dispersions estimated by the estimateDisp function from edgeR (McCarthy et al., 2012; Robinson et al., 2010) and a false discovery rate (FDR) cut-off of 0.05.

### 10× Single Cell RNAseq and Data Analysis

Data were processed by scPipe, with reads mapped to the mm10 mouse genome using the Subread aligner. The low-quality cells were filtered out with detect_outlier function from scPipe and cells with less than 1000 UMI counts or 800 detected genes being removed. After quality control, the gene count matrix was used as input to the Seurat pipeline for downstream analysis with default parameters (Butler et al., 2018). Cell cycle effect was removed using Seurat with the ScaleData function. The first 10 principal components from PCA were used as input for Seurat clustering algorithm, with default resolution 0.8. The pooled samples were demultiplexed by the expression of sex specific genes DDX3Y and XIST. The GO term enrichment analysis was performed by topGO with default parameters.

### Live-Cell Metabolic Assays

Cellular bioenergetics were measured utilizing the Seahorse XF24-3 Bioanalyzer according to manufacturer’s instructions (Agilent Technologies, Santa Clara, California, USA). Briefly, MuSCs (or C2C12 cells) were seeded into a 24-well microplate suspended in 200 µL of media containing either HG-, LG-or GAL-based growth media and allowed to attach overnight. At least three wells were left blank to allow for background correction. One hour prior to the commencement of the assay, cells were washed with XF base media (HCO3- free modified DMEM, Agilent, Santa Clara, California, USA) supplemented with 1 mM sodium pyruvate (Gibco), 1 mM L-glutamine (Gibco) and 25 mM glucose (Sigma-Aldrich) and resuspended in 630 µL of XF base media. Cells were incubated for one hour at 37°C in a CO_2_ free incubator to allow equilibration. Basal metabolism was then assessed by concurrently measuring the oxygen consumption rate (OCR) and extracellular acidification rate (ECAR). Following measurements of basal metabolism, a mitochondrial stress test was performed as described previously (Nicholls et al., 2010).

### Flow Cytometry

Primary mouse MuSCs were seeded in 6 well plates at a density of 5 × 10^4^ cells/well and cultured in either HG-, LG-or GAL-based growth media for 96 hrs. Cells were then cultured for an additional one hour in growth media containing 100 nM Mitotracker Red CMXRos. Cells were subsequently trypsinized and centrifuged at 1600 rpm for 5 mins. The cell pellet was washed with PBS and fixed in 1 mL ice cold 4% PFA (Electron Microscopy Science, Hatfield, PA, USA) diluted in PBS. Flow cytometric analyses were performed on a BD Fortessa 4-Laser Analyzer (Beckman Coulter, Mount Waverley, VIC, Australia).

### Western Blot Analyses

Whole cell lysates were prepared from cells following culture in either HG-, LG-or GAL-based growth media. Cells were washed with PBS before being incubated in Trypsin-EDTA (0.25%, Thermo Scientific, Waltham, Massachusetts, USA) to detach cells from cell culture plates. Cells were centrifuged at 1,600 rpm for 5 min, with the resulting pellet washed twice in ice cold PBS before being snap frozen in liquid nitrogen and stored at -80°C. Prior to lysing, pellets were washed with ice cold PBS containing 1 mM phenylmethylsulfonyl fluoride (PMSF, Pierce Biotechnology, Rockford, Illinois, USA), 0.1% protease inhibitor cocktail (P8340, Sigma Aldrich) and 0.1% phosphatase inhibitor cocktail (P2850, P5726, Sigma Aldrich). Cells were lysed in ice cold radio-immunoprecipitation assay (RIPA) lysis buffer (50 mM TrisHCL, 150 mM NaCl, 1 mM EDTA, 1% Triton X-100, 1% Na-Deoxycholic acid, 0.1% SDS). To ensure complete breakdown of cell and nuclear membranes, cells were sonicated for 10 s (Microson Ultrasonic Cell Disruptor, Misonix, Farmingdale, New York, USA).

Cell lysates were centrifuged at 10,000 g for 10 min and the supernatant harvested. The cell lysate protein concentration was measured via a standard Lowry protein assay and then equalized to a concentration of 1 µg/µL. Samples were denatured at 95°C for 5 min and equal amounts of protein (20 µg diluted in 4× Laemmli loading buffer) was loaded into precast sodium dodecyl sulphate (SDS)-polyacrylamide gels. Proteins were then separated based on their electrophoretic mobility by exposing the polyacrylamide gel to a constant voltage of 150V, until the migrating dye front reached the bottom of the gel. Equal protein loading was confirmed using a stain-free visualization system (Criterion TGX Stain-Free Precast Gels, BioRad Laboratories, Hatfield, PA, USA) and the protein transferred to 0.45 mm polyvinylidene difluoride (PVDF) membrane using the Trans-Blot Turbo transfer system (BioRad Laboratories, Hatfield, PA, USA).

Non-specific binding sites were blocked in 3% Bovine serum albumen (BSA) in Tris-buffered saline Tween 20 (TBST) at room temperature for two hours. Membranes were incubated overnight in primary antibodies at 4°C made up in blocking buffer. Membranes were then washed in TBST before being incubated for 2 hours at room temperature in horseradish peroxidase-conjugated secondary antibodies diluted in blocking buffer. Following 6 × 10 min washes in TBST, membranes were labelled with Supersignal West Femto Chemiluminescent Substrate (Pierce Biotechnology, Rockford, Illinois, USA) and imaged using the ChemiDoc XRS (Biorad, Hercules, CA, USA). The final protein bands were quantified using ImageLab Software (Biorad, Hercules, CA, USA).

### Metabolite Assays

Colorimetric and fluorometric assays were performed as directed by the manufacturer’s instructions to determine the concentration of key metabolites; PEP, pyruvate, acetyl-CoA, citrate and α-ketoglutarate (Biovision Inc., Milipitas, CA, USA). Briefly, C2C12 cells were cultured for 24 hrs in either HG-, LG-or GAL-based growth media before being trypsinized and snap frozen in liquid nitrogen. 1-5×10^6^ cells were lysed and then centrifuged at 10,000 rpm at 4°C to remove any insoluble material. Cell lysates were incubated in the presence of a colorimetric/fluorometric probe at room temperature for 30 mins and the absorbance read on a photospectrometer (Multiskan, Thermo Scientific, Waltham, MA, USA) at 570 nm (for colorimetric assays) and 535 nm (for fluorometric assays). Metabolite concentrations were then extrapolated from a standard curve and corrected to total protein within the cell lysates.

### Immunofluorescence

Cells (or fibers) were fixed with 4% paraformaldehyde (PFA, Electron Microscopy Science, Hatfield, PA, USA) diluted in PBS for 15 mins and then permeabilized with 0.1% Triton X-100 (PBST, BioRad Laboratories, Hatfield, PA, USA) in PBS for 10 min. Cells (or fibers) were then blocked with 3% bovine serum albumin (BSA, Sigma-Aldrich, St. Louis, MO, USA) diluted in PBST for 1 hr at room temperature to block non-specific binding sites. Cells (or fibers) were then incubated overnight in blocking solution containing desired primary antibodies. The following day, cells (or fibers) were then incubated with the appropriate secondary antibody diluted 1:1000 in blocking medium for 2 hr at room temperature. Following this incubation, the cells (or fibers) were washed with PBS and counterstained with 4’6- diamidino-2-phenylindole (DAPI, 5 mg/ml, Molecular Probes, Invitrogen, Carlsbad, USA) in PBS to label nuclei.

### Myofiber Isolation and Culture

Extensor digitorum longus (EDL) muscles were isolated from the hindlimbs of 12 wk old male C57BL/6 mice, and incubated in 0.2% collagenase (type I) diluted in DMEM) at 37°C for 45 min, as described previously (Pasut et al., 2013). At the completion of this incubation, EDL muscles were transferred to growth media containing DMEM (cat# 11966025, Gibco) supplemented with 1 mM sodium pyruvate (cat# 11360070, Gibco), 10% fetal bovine serum (FBS, Australian origin, cat# 10099141, Gibco), 10% horse serum (HS, New Zealand origin, cat# 16050130, Gibco), 0.05% chick embryo extract (cat# C3999, US Biological), and 1% penicillin/streptomycin (Gibco). Individual myofibers were liberated from the EDL via gentle agitation with growth media, and then individually plated in 48-well plates in growth media supplemented with either 25 mM or 5 mM glucose, or 10 mM galactose.

## QUANTIFICATION AND STATISTICAL ANALYSIS

Statistical analyses of data were performed using GraphPad Prism software (GraphPad Software, CA, USA). All values are expressed as mean ± standard error of the mean (SEM) unless otherwise stated. One-way and two-way ANOVA were used to compare differences between groups with Fisher’s LSD post hoc test used to test for significant differences between groups. Where data did not conform to a normal distribution, results are reported as mean and 95% confidence interval (CI) with results considered significant where there was no overlap of CI. Significance was set at *p<0.05 and **p<0.01. All analyses of data concerning immunofluorescence images were performed in a double-blinded manner.

## DATA AND SOFTWARE AVAILABILITY

All software used for the analysis of scRNAseq or population RNAseq is freely available as indicated in the STAR methods table. Both MuSC scRNAseq and whole population RNAseq data have been deposited at NCBI Gene Expression Omnibus (Accession numbers GSE154868 (short-read data only), GSE117386 and GSE117229 respectively). MALDI-MSI datasets are freely available for download from www.metaspace2020.eu with the following names: muscle_01_rms and muscle_02_rms, corresponding to the control and treated tissue respectively.

## SUPPLEMENTAL INFORMATION

Supplemental information includes, four figures, and five tables.

## ACKNOWLEDGMENTS

We thank Dr Vanta Jameson for help in cell sorting and flow-cytometry. The authors would like to thank all members of the Centre for Muscle Research in the Department of Anatomy and Physiology at The University of Melbourne for useful discussion and feedback. CHL was supported by an Australian Postgraduate Award from the Australian Government. This work was supported by project grants from the Australian Research Council (DP150100206) and the National Health and Medical Research Council (GNT1120714) awarded to JGR and GSL.

## AUTHOR CONTRIBUTIONS

CHL and JGR designed the experiments. CHL, GSL and JGR wrote the manuscript. CHL performed most of the presented experiments. JT helped with FACS, immunoblotting and cell culture, DZ and SN performed scRNAseq, SS, LT, SN and MR conducted downstream bioinformatics of scRNAseq, BAB conducted MALDI-MSI and data analysis. JGR performed RNAseq and downstream bioinformatics of C2C12 cells. Funding to complete the experiments was obtained by JGR and GSL.

## Supplemental information

**Figure S1.**
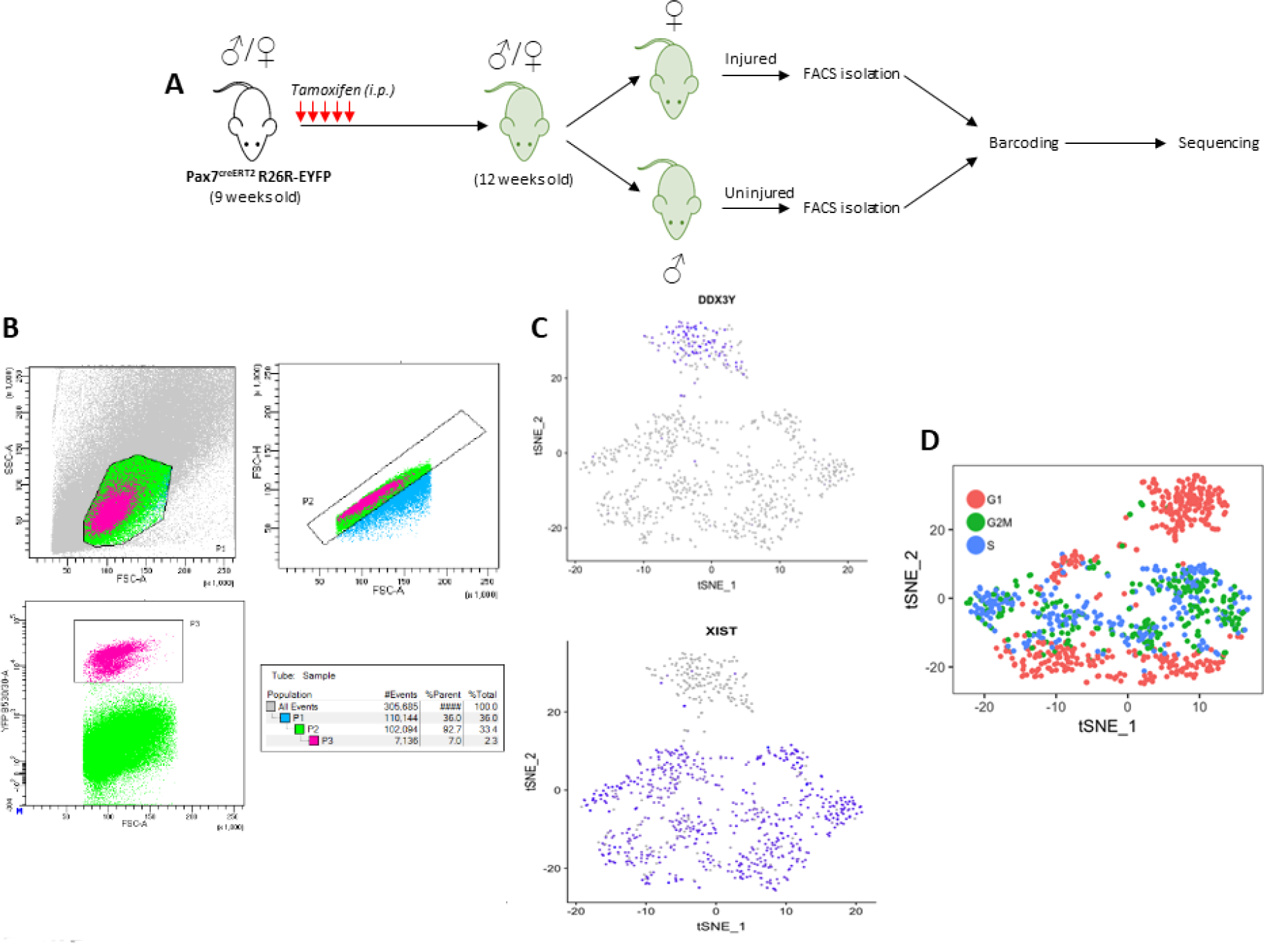
Related to Figure 2. Isolation of Primary MuSCs and Single Cell RNAseq. (A) To rapidly and efficiently isolate large numbers of primary MuSCs for scRNAseq, we utilized the Pax7^creERT2^R26-EYFP mouse. A schematic showing the timeline for tamoxifen injection, muscle injury and the isolation of primary MuSCs is shown. The TA muscles of female mice were injected with BaCl_2_ to induce muscle injury, while the TA muscles of male mice served as uninjured control muscles. (B) FACS gating strategies for the isolation of YFP^+^ primary MuSCs (Based on Liu et al., 2015). An individual mouse yielded between 200,000 to 250,000 YFP^+^ cells. (C) X (*Xist*) and Y (*Ddx3y*) chromosome specific genes were used to identify MuSCs isolated from injured (female) and uninjured (male) muscles. (D) MuSCs isolated from uninjured muscles were confirmed to be in the G0/G1 phases of the cell cycle.

**Figure S2.**
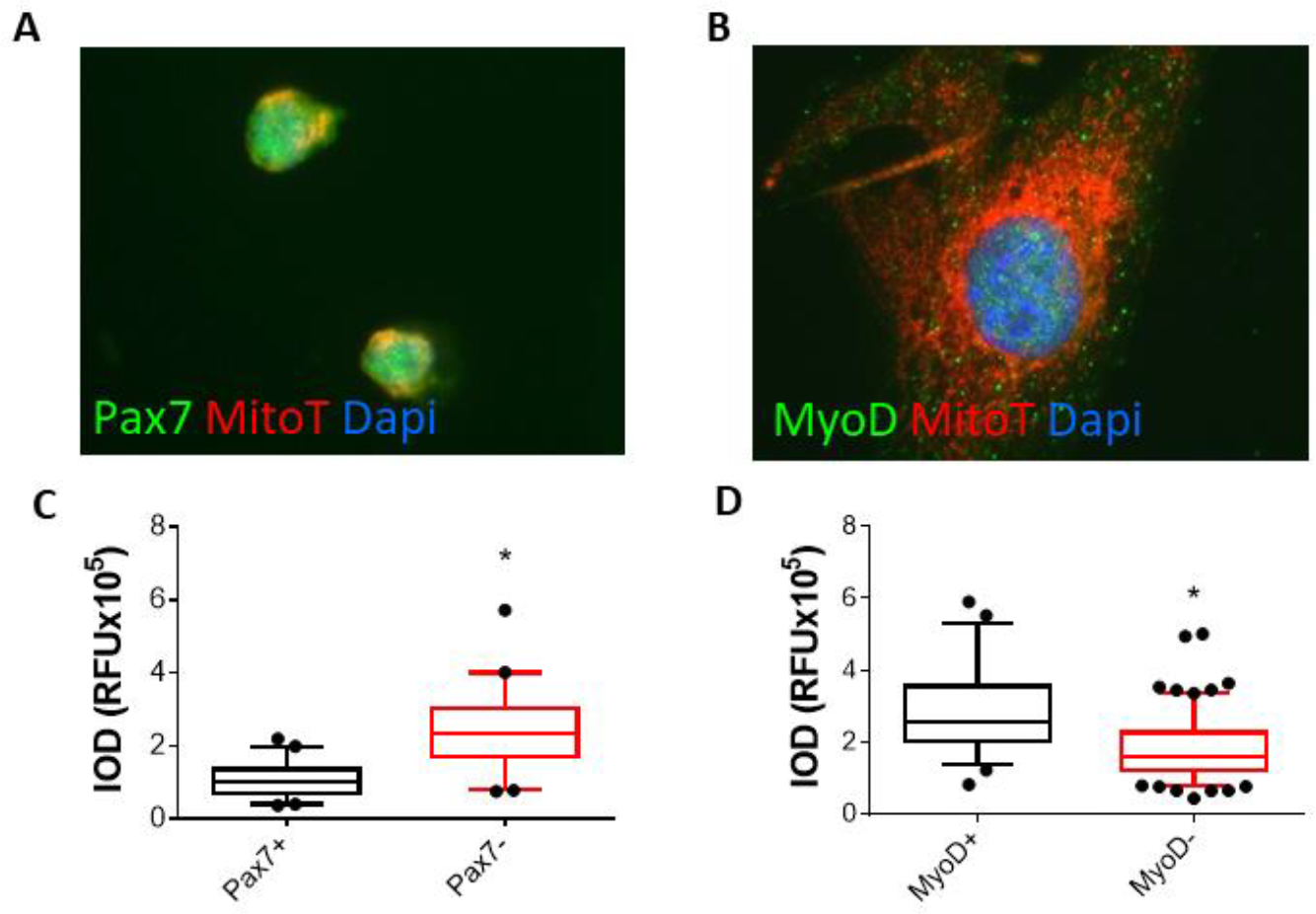
Related to Figure 2. Myogenic Specification and Proliferation is Linked to Cellular Metabolism. (A,B) To examine the link between myogenic specification and mitochondrial content we labeled proliferating C2C12 cells with MitoTracker Red and either Pax7 (A) or MyoD (B). (C) Quantification of the IOD of MitoTracker labeling in Pax7^+^ and Pax7^-^ cells. (C) Quantification of the IOD of MitoTracker labeling in MyoD^+^ and MyoD^-^ cells. Student’s t-test, *p<0.05

**Figure S3.**
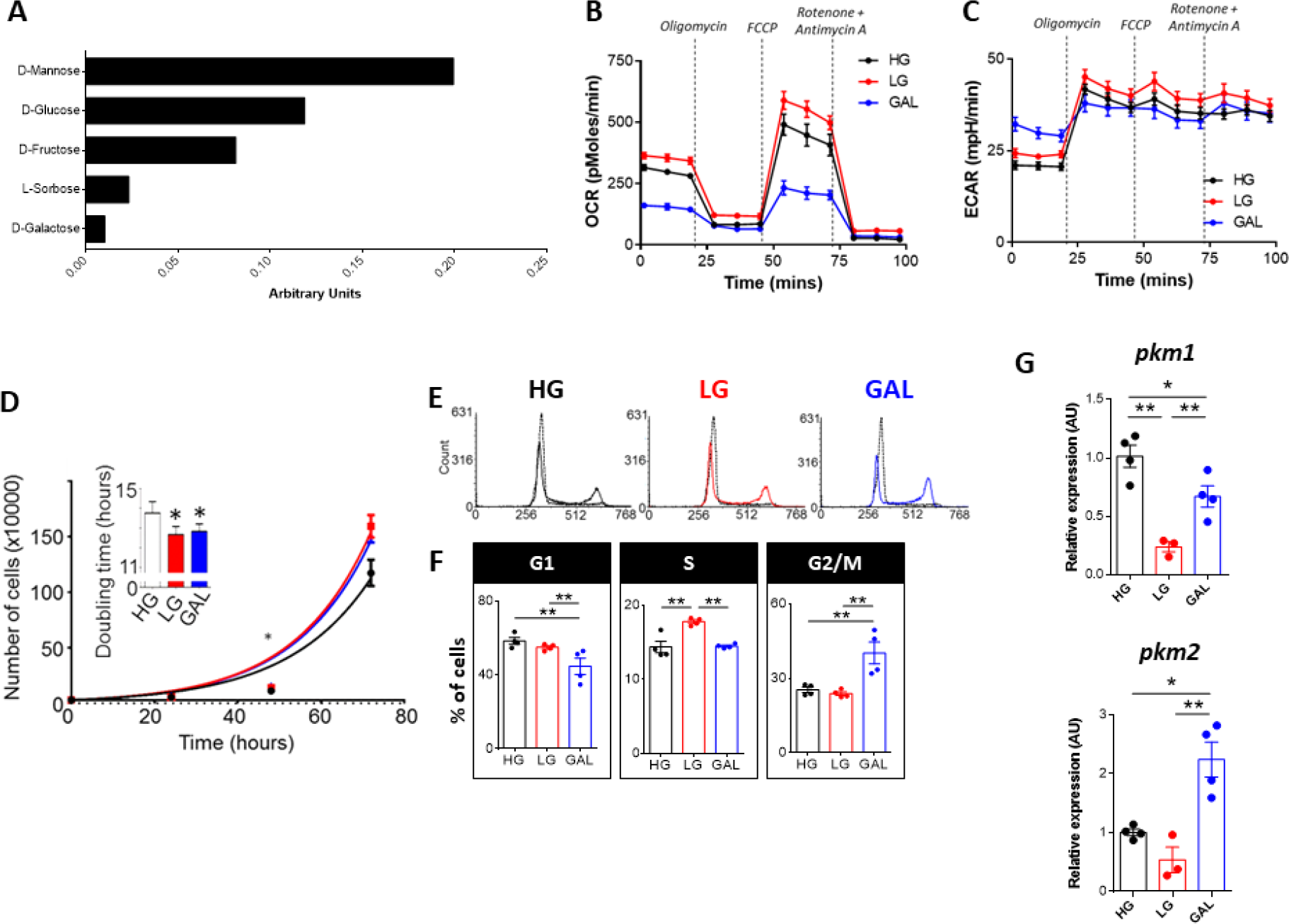
Related to Figure 3. Monosaccharide Availability in the Extracellular Environment Regulates Innate Metabolism and Proliferation of C2C12 Cells. (A) Proliferating C2C12 cells were cultured in the presence of different monosaccharides to determine substrate utilization, using a commercially available microarray for carbon-energy utilization (Biolog). (B) A mitochondrial stress test was performed on C2C12 cells cultured for 24 hrs in high glucose (HG), low glucose (LG), or galactose (GAL) based growth media (n=6-7/group, mean ± SEM). (C) The extracellular acidification rate (ECAR) as measured during (B). (D) C2C12 cells were seeded in HG-, LG-or GAL-based growth media and total cell numbers were determined at 24, 48, and 72 hrs post-seeding. The mean doubling time was then calculated from the resulting graph (n=4/group, mean ± SEM). (E-F) Propidium iodide labelling coupled with flow cytometric analyses revealed an increased proportion of LG cultured cells in S-phase and an increased proportion of GAL cultured cells in G2/M-phase. (G) Differential splicing of *pkm* following 24 hrs of culture in either HG, LG, or GAL was determined by qPCR (n=3-4 replicates/group). ANOVA with Fisher’s LSD multiple comparison procedure, *p<0.05, **p<0.01

**Figure S4.**
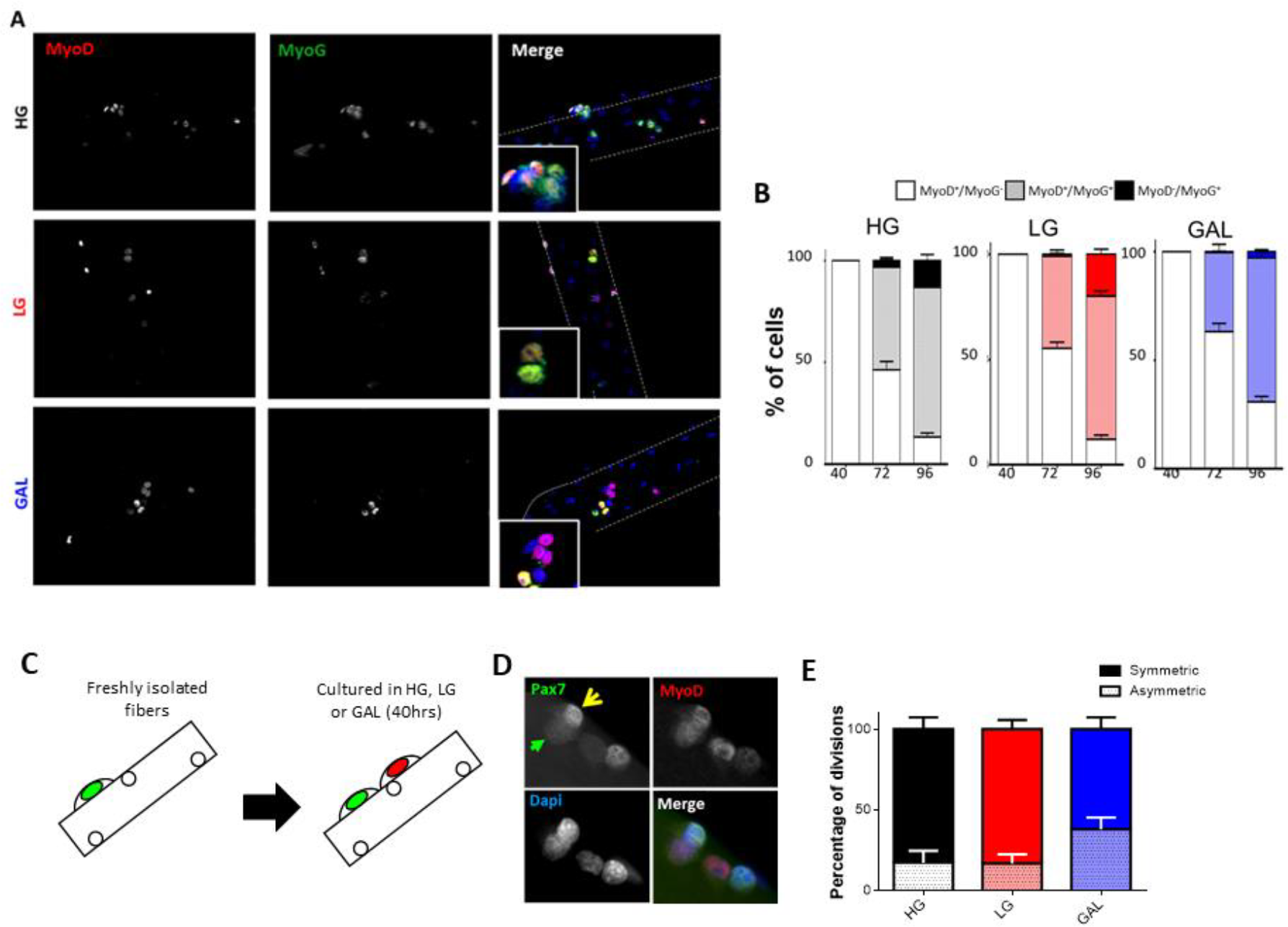
Related to Figure 7. Monosaccharide Availability in the Extracellular Environment Regulates Differentiation of MuSCs. (A) Single muscle fibers were cultured in HG, LG, or GAL for 40, 72, or 96 hrs and immunofluorescently labelled with MyoD and MyoG. (B) Quantification of the proportion of activated MuSCs/committed muscle progenitors (MyoD^+^/MyoG^-^), early differentiated progenitors (MyoD^+^/MyoG^+^), and late differentiated progenitors (MyoD^-^/MyoG^+^), on fibers cultured for 40, 72, or 96 hrs in HG, LG, or GAL (n>80 fibers/timepoint, isolated from n=6 mice, mean ± SEM). (C) Schematic detailing the analysis of MuSC asymmetric division at 40 hrs of culture. (D) Single muscle fibers were cultured in HG, LG, or GAL for 40 hrs (approximate time of first division) and immunofluorescently labelled with Pax7 and MyoD to identify symmetrically and asymmetrically dividing MuSCs. (E) Quantification of (E). (n>80 fibers, isolated from n=6 mice, mean ± SEM).

**Table S1. Related to Figure 1. Differential Metabolite Expression from Metabolomics Performed on Uninjured and Injured Skeletal Muscles**

**Table S2. Related to Figure 2. Differential Gene Expression from scRNAseq Performed on Freshly Isolated MuSCs from Uninjured and Injured Skeletal Muscles**

**Table S3. Related to Figure 5. Differential Gene Expression from Population RNAseq Performed on Primary MuSCs either Freshly Isolated, or Following 24, 48, or 72 hrs of Culture in Growth Media**

**Table S4. Related to Figure 5. Differential Gene Expression from Population RNAseq Performed on Primary MuSCs Cultured for 48 hrs in Growth Media Supplemented with HG, LG, or GAL.**

**Table S5. Related to Figure 6. Differential Gene Expression from scRNAseq Performed on Freshly Isolated and MuSCs cultured for 96 hrs in HG, LG, or GAL**

## REFERENCES

Butler, A., Hoffman, P., Smibert, P., Papalexi, E., and Satija, R. (2018). Integrating single-cell transcriptomic data across different conditions, technologies, and species. Nature biotechnology 36, 411–420.

Cai, L., Sutter, B.M., Li, B., and Tu, B.P. (2011). Acetyl-CoA induces cell growth and proliferation by promoting the acetylation of histones at growth genes. Molecular cell 42, 426–437.

Casella, J.F., Flanagan, M.D., and Lin, S. (1981). Cytochalasin D inhibits actin polymerization and induces depolymerization of actin filaments formed during platelet shape change. Nature 293, 302–305.

Cerletti, M., Jang, Y.C., Finley, L.W., Haigis, M.C., and Wagers, A.J. (2012). Short-term calorie restriction enhances skeletal muscle stem cell function. Cell Stem Cell 10, 515–519.

Chang, N.C., Sincennes, M.C., Chevalier, F.P., Brun, C.E., Lacaria, M., Segales, J., Munoz-Canoves, P., Ming, H., and Rudnicki, M.A. (2018). The Dystrophin Glycoprotein Complex Regulates the Epigenetic Activation of Muscle Stem Cell Commitment. Cell stem cell 22, 755–768.e756.

Cheung, T.H., Quach, N.L., Charville, G.W., Liu, L., Park, L., Edalati, A., Yoo, B., Hoang, P., and Rando, T.A. (2012). Maintenance of muscle stem-cell quiescence by microRNA-489. Nature 482, 524–528.

Cox, A.G., Tsomides, A., Yimlamai, D., Hwang, K.L., Miesfeld, J., Galli, G.G., Fowl, B.H., Fort, M., Ma, K.Y., Sullivan, M.R., et al. (2018). Yap regulates glucose utilization and sustains nucleotide synthesis to enable organ growth. The EMBO journal 37.

Dell’Orso, S., Juan, A.H., Ko, K.D., Naz, F., Perovanovic, J., Gutierrez-Cruz, G., Feng, X., and Sartorelli, V. (2019). Single cell analysis of adult mouse skeletal muscle stem cells in homeostatic and regenerative conditions. Development (Cambridge, England) 146.

Dumont, N.A., Wang, Y.X., von Maltzahn, J., Pasut, A., Bentzinger, C.F., Brun, C.E., and Rudnicki, M.A. (2015). Dystrophin expression in muscle stem cells regulates their polarity and asymmetric division. Nat Med 21, 1455–1463.

Fukada, S., Uezumi, A., Ikemoto, M., Masuda, S., Segawa, M., Tanimura, N., Yamamoto, H., Miyagoe-Suzuki, Y., and Takeda, S. (2007). Molecular signature of quiescent satellite cells in adult skeletal muscle. Stem cells (Dayton, Ohio) 25, 2448–2459.

Gatta, L., Vitiello, L., Gorini, S., Chiandotto, S., Costelli, P., Giammarioli, A.M., Malorni, W., Rosano, G., and Ferraro, E. (2017). Modulating the metabolism by trimetazidine enhances myoblast differentiation and promotes myogenesis in cachectic tumor-bearing c26 mice. Oncotarget 8, 113938–113956.

Hardy, D., Besnard, A., Latil, M., Jouvion, G., Briand, D., Thépenier, C., Pascal, Q., Guguin, A., Gayraud-Morel, B., Cavaillon, J.M., et al. (2016). Comparative Study of Injury Models for Studying Muscle Regeneration in Mice. PloS one 11, e0147198.

Hashimshony, T., Senderovich, N., Avital, G., Klochendler, A., de Leeuw, Y., Anavy, L., Gennert, D., Li, S., Livak, K.J., Rozenblatt-Rosen, O., et al. (2016). CEL-Seq2: sensitive highly-multiplexed single-cell RNA-Seq. Genome biology 17, 77.

Hernandez-Hernandez, J.M., Garcia-Gonzalez, E.G., Brun, C.E., and Rudnicki, M.A. (2017). The myogenic regulatory factors, determinants of muscle development, cell identity and regeneration. Seminars in cell & developmental biology 72, 10–18.

Hosios, A.M., Fiske, B.P., Gui, D.Y., and Vander Heiden, M.G. (2015). Lack of Evidence for PKM2 Protein Kinase Activity. Molecular cell 59, 850–857.

Hosios, A.M., Hecht, V.C., Danai, L.V., Johnson, M.O., Rathmell, J.C., Steinhauser, M.L., Manalis, S.R., and Vander Heiden, M.G. (2016). Amino Acids Rather than Glucose Account for the Majority of Cell Mass in Proliferating Mammalian Cells. Developmental cell 36, 540–549.

Johnson, W.E., Li, C., and Rabinovic, A. (2007). Adjusting batch effects in microarray expression data using empirical Bayes methods. Biostatistics (Oxford, England) 8, 118–127.

Kuang, S., Kuroda, K., Le Grand, F., and Rudnicki, M.A. (2007). Asymmetric self-renewal and commitment of satellite stem cells in muscle. Cell 129, 999–1010.

Liao, Y., Smyth, G.K., and Shi, W. (2013). The Subread aligner: fast, accurate and scalable read mapping by seed-and-vote. Nucleic acids research 41, e108.

Liu, L., Cheung, T.H., Charville, G.W., and Rando, T.A. (2015). Isolation of skeletal muscle stem cells by fluorescence-activated cell sorting. Nature protocols 10, 1612–1624.

Lun, A.T., McCarthy, D.J., and Marioni, J.C. (2016). A step-by-step workflow for low-level analysis of single-cell RNA-seq data with Bioconductor. F1000Research 5, 2122.

Lunt, S.Y., Muralidhar, V., Hosios, A.M., Israelsen, W.J., Gui, D.Y., Newhouse, L., Ogrodzinski, M., Hecht, V., Xu, K., Acevedo, P.N., et al. (2015). Pyruvate kinase isoform expression alters nucleotide synthesis to impact cell proliferation. Molecular cell 57, 95–107.

Lunt, S.Y., and Vander Heiden, M.G. (2011). Aerobic glycolysis: meeting the metabolic requirements of cell proliferation. Annual review of cell and developmental biology 27, 441–464.

Ly, C.H., Lynch, G.S., and Ryall, J.G. (2020). A Metabolic Roadmap for Somatic Stem Cell Fate. Cell metabolism 31, 1052–1067.

Machado, L., Esteves de Lima, J., Fabre, O., Proux, C., Legendre, R., Szegedi, A., Varet, H., Ingerslev, L.R., Barrès, R., Relaix, F., et al. (2017). In Situ Fixation Redefines Quiescence and Early Activation of Skeletal Muscle Stem Cells. Cell reports 21, 1982–1993.

McCarthy, D.J., Chen, Y., and Smyth, G.K. (2012). Differential expression analysis of multifactor RNA-Seq experiments with respect to biological variation. Nucleic acids research 40, 4288–4297.

Murphy, M.M., Lawson, J.A., Mathew, S.J., Hutcheson, D.A., and Kardon, G. (2011). Satellite cells, connective tissue fibroblasts and their interactions are crucial for muscle regeneration. Development (Cambridge, England) 138, 3625-3637.

Nguyen, J.H., Chung, J.D., Lynch, G.S., and Ryall, J.G. (2019). The Microenvironment Is a Critical Regulator of Muscle Stem Cell Activation and Proliferation. Frontiers in cell and developmental biology 7, 254.

Nicholls, D.G., Darley-Usmar, V.M., Wu, M., Jensen, P.B., Rogers, G.W., and Ferrick, D.A. (2010). Bioenergetic profile experiment using C2C12 myoblast cells. Journal of visualized experiments : JoVE.

Olguin, H.C., Yang, Z., Tapscott, S.J., and Olwin, B.B. (2007). Reciprocal inhibition between Pax7 and muscle regulatory factors modulates myogenic cell fate determination. J Cell Biol 177, 769–779.

Pala, F., Di Girolamo, D., Mella, S., Yennek, S., Chatre, L., Ricchetti, M., and Tajbakhsh, S. (2018). Distinct metabolic states govern skeletal muscle stem cell fates during prenatal and postnatal myogenesis. Journal of cell science 131.

Pallafacchina, G., François, S., Regnault, B., Czarny, B., Dive, V., Cumano, A., Montarras, D., and Buckingham, M. (2010). An adult tissue-specific stem cell in its niche: a gene profiling analysis of in vivo quiescent and activated muscle satellite cells. Stem cell research 4, 77–91.

Pasut, A., Jones, A.E., and Rudnicki, M.A. (2013). Isolation and culture of individual myofibers and their satellite cells from adult skeletal muscle. Journal of visualized experiments : JoVE, e50074.

Ritchie, M.E., Phipson, B., Wu, D., Hu, Y., Law, C.W., Shi, W., and Smyth, G.K. (2015). limma powers differential expression analyses for RNA-sequencing and microarray studies. Nucleic acids research 43, e47.

Robinson, M.D., McCarthy, D.J., and Smyth, G.K. (2010). edgeR: a Bioconductor package for differential expression analysis of digital gene expression data. Bioinformatics (Oxford, England) 26, 139–140.

Rocheteau, P., Gayraud-Morel, B., Siegl-Cachedenier, I., Blasco, M.A., and Tajbakhsh, S. (2012). A subpopulation of adult skeletal muscle stem cells retains all template DNA strands after cell division. Cell 148, 112–125.

Rodgers, J.T., King, K.Y., Brett, J.O., Cromie, M.J., Charville, G.W., Maguire, K.K., Brunson, C., Mastey, N., Liu, L., Tsai, C.R., et al. (2014). mTORC1 controls the adaptive transition of quiescent stem cells from G0 to G(Alert). Nature 510, 393–396.

Ryall, J.G., Cliff, T., Dalton, S., and Sartorelli, V. (2015a). Metabolic Reprogramming of Stem Cell Epigenetics. Cell Stem Cell 17, 651–662.

Ryall, J.G., Dell’Orso, S., Derfoul, A., Juan, A., Zare, H., Feng, X., Clermont, D., Koulnis, M., Gutierrez-Cruz, G., Fulco, M., et al. (2015b). The NAD(+)-dependent SIRT1 deacetylase translates a metabolic switch into regulatory epigenetics in skeletal muscle stem cells. Cell Stem Cell 16, 171–183.

Shang, M., Cappellesso, F., Amorim, R., Serneels, J., Virga, F., Eelen, G., Carobbio, S., Rincon, M.Y., Maechler, P., De Bock, K., et al. (2020). Macrophage-derived glutamine boosts satellite cells and muscle regeneration. Nature 587, 626–631.

Shea, K.L., Xiang, W., LaPorta, V.S., Licht, J.D., Keller, C., Basson, M.A., and Brack, A.S. (2010). Sprouty1 regulates reversible quiescence of a self-renewing adult muscle stem cell pool during regeneration. Cell stem cell 6, 117–129.

Siegel, A.L., Atchison, K., Fisher, K.E., Davis, G.E., and Cornelison, D.D. (2009). 3D timelapse analysis of muscle satellite cell motility. Stem cells (Dayton, Ohio) 27, 2527–2538.

Srinivas, S., Watanabe, T., Lin, C.S., William, C.M., Tanabe, Y., Jessell, T.M., and Costantini, F. (2001). Cre reporter strains produced by targeted insertion of EYFP and ECFP into the ROSA26 locus. BMC developmental biology 1, 4.

Su, S., Law, C.W., Ah-Cann, C., Asselin-Labat, M.L., Blewitt, M.E., and Ritchie, M.E. (2017). Glimma: interactive graphics for gene expression analysis. Bioinformatics (Oxford, England) 33, 2050–2052.

Theret, M., Gsaier, L., Schaffer, B., Juban, G., Ben Larbi, S., Weiss-Gayet, M., Bultot, L., Collodet, C., Foretz, M., Desplanches, D., et al. (2017). AMPKalpha1-LDH pathway regulates muscle stem cell self-renewal by controlling metabolic homeostasis. The EMBO journal 36, 1946–1962.

Tian, L., Su, S., Dong, X., Amann-Zalcenstein, D., Biben, C., Seidi, A., Hilton, D.J., Naik, S.H., and Ritchie, M.E. (2018). scPipe: A flexible R/Bioconductor preprocessing pipeline for single-cell RNA-sequencing data. PLoS computational biology 14, e1006361.

Wellen, K.E., Hatzivassiliou, G., Sachdeva, U.M., Bui, T.V., Cross, J.R., and Thompson, C.B. (2009). ATP-citrate lyase links cellular metabolism to histone acetylation. Science 324, 1076–1080.

Yucel, N., Wang, Y.X., Mai, T., Porpiglia, E., Lund, P.J., Markov, G., Garcia, B.A., Bendall, S.C., Angelo, M., and Blau, H.M. (2019). Glucose Metabolism Drives Histone Acetylation Landscape Transitions that Dictate Muscle Stem Cell Function. Cell reports 27, 3939–3955.e3936.

Zammit, P.S. (2017). Function of the myogenic regulatory factors Myf5, MyoD, Myogenin and MRF4 in skeletal muscle, satellite cells and regenerative myogenesis. Seminars in cell & developmental biology 72, 19–32.

Zhang, J., Muri, J., Fitzgerald, G., Gorski, T., Gianni-Barrera, R., Masschelein, E., D’Hulst, G., Gilardoni, P., Turiel, G., Fan, Z., et al. (2020). Endothelial Lactate Controls Muscle Regeneration from Ischemia by Inducing M2-like Macrophage Polarization. Cell metabolism 31, 1136–1153.e1137.

Zhao, S., Torres, A., Henry, R.A., Trefely, S., Wallace, M., Lee, J.V., Carrer, A., Sengupta, A., Campbell, S.L., Kuo, Y.M., et al. (2016). ATP-Citrate Lyase Controls a Glucose-to-Acetate Metabolic Switch. Cell reports 17, 1037–1052.

